# High-performing neural network models of visual cortex benefit from high latent dimensionality

**DOI:** 10.1101/2022.07.13.499969

**Authors:** Eric Elmoznino, Michael F. Bonner

## Abstract

Geometric descriptions of deep neural networks (DNNs) have the potential to uncover core representational principles of computational models in neuroscience. Here we examined the geometry of DNN models of visual cortex by quantifying the latent dimensionality of their natural image representations. A popular view holds that optimal DNNs compress their representations onto low-dimensional subspaces to achieve invariance and robustness, which suggests that better models of visual cortex should have lower dimensional geometries. Surprisingly, we found a strong trend in the opposite direction—neural networks with high-dimensional image subspaces tended to have better generalization performance when predicting cortical responses to held-out stimuli in both monkey electrophysiology and human fMRI data. Moreover, we found that high dimensionality was associated with better performance when learning new categories of stimuli, suggesting that higher dimensional representations are better suited to generalize beyond their training domains. These findings suggest a general principle whereby high-dimensional geometry confers computational benefits to DNN models of visual cortex.

## 1 Introduction

Deep neural networks (DNNs) are the predominant framework for computational modeling in neuroscience [73, 92, 43, 52, 59]. When using DNNs to model neural systems, one of the fundamental questions that researchers hope to answer is: What core factors explain why some DNNs succeed and others fail? Researchers often attribute the success of DNNs to explicit design choices in a model’s construction, such as its architecture, learning objective, and training data [92, 13, 14, 52, 59, 93, 48, 22, 50, 97, 12, 20]. However, an alternative perspective explains DNNs through the geometry of their latent representational subspaces, which abstracts over the details of how networks are constructed [19, 85, 17, 16, 47]. Here we sought to understand the geometric principles that underlie the performance of DNN models of visual cortex.

We examined the geometry of DNNs by quantifying the dimensionality of their representational subspaces. DNN models of vision contain thousands of artificial neurons, but their representations are known to be constrained to lower-dimensional subspaces that are embedded within the ambient space of the neural population [e.g. 3]. Many have argued that DNNs benefit from representing stimuli in subspaces that are as low-dimensional as possible, and it is proposed that low dimensionality improves a network’s generalization performance, its robustness to noise, and its ability to separate stimuli into meaningful categories [3, 72, 23, 25, 24, 57, 70, 96, 90, 38, 2, 61, 49]. Similar arguments have been made for the benefits of low-dimensional subspaces in the sensory, motor, and cognitive systems of the brain [18, 33, 67, 64, 58, 31, 77]. However, contrary to this view, there are also potential benefits of high-dimensional subspaces, including the efficient utilization of a network’s representational resources and increased expressivity, making for a greater number of potential linear readouts [88, 81, 4, 30, 55, 66].

We wondered whether the dimensionality of representational subspaces might be relevant for understanding the relationship between DNNs and visual cortex and, if so, what level of dimensionality performs best. To answer these questions, we measured the latent dimensionality of DNNs trained on a variety of supervised and self-supervised tasks using multiple datasets, and we assessed their accuracy at predicting image-evoked activity patterns in visual cortex for held-out stimuli using both monkey electrophysiology and human fMRI data. We discovered a powerful relationship between dimensionality and accuracy: specifically, we found that DNNs with higher latent dimensionality explain more variance in the image representations of high-level visual cortex. This was true even when controlling for model depth and the number of parameters in each network, and it could not be explained by overfitting because our analyses explicitly tested each network’s ability to generalize to held-out stimuli. Furthermore, we found that high dimensionality also conferred computational benefits when learning to classify new categories of stimuli, providing support for its adaptive role in visual behaviors. Together, these findings suggest that high-performing computational models of visual cortex are characterized by high-dimensional representational subspaces, allowing them to efficiently support a greater diversity of linear readouts for natural images.

## 2 Results

### 2.1 Dimensionality and alignment in computational brain models

We set out to answer two fundamental questions about the geometry of DNNs in computational neuroscience. First, is there a relationship between latent dimensionality and DNN performance? Second, if latent dimensionality is indeed related to DNN performance, what level of dimensionality is better? In other words, do DNN models of neural systems primarily benefit from the robustness and invariance of low-dimensional codes or the expressivity of high-dimensional codes? To explore the theoretical issues underlying these questions, we first performed simulations that illustrate how the geometry of latent subspaces might influence the ability of representational models to account for variance in brain activity patterns.

For our simulations, we considered a scenario in which all brain representations and all relevant computational models are sampled from a large subspace of image representations called the *natural image subspace*. Here, we use the term subspace to describe the lower-dimensional subspace spanned by the major principal components of a system with higher ambient dimensionality (e.g., neurons). We sampled observations from this natural image subspace and projected these observations onto the dimensions spanned by two smaller subspaces called the *ecological subspace* and the *model subspace*. Projections onto the ecological subspace simulate image representations in the brain, and projections onto the model subspace simulate image representations in a computational model. We analyzed these simulated data using a standard approach in computational neuroscience, known as the encoding-model approach. Specifically, we mapped model representations to brain representations using cross-validated linear regression. This analysis yielded an encoding score, which is the explained variance for held-out data in the cross-validated regression procedure. Computational models with higher encoding scores have better performance when predicting brain representations for held-out data. Further details regarding the theoretical grounding and technical implementation of our simulations are provided in Appendices B and C.

Using this simulation framework, we can now illustrate how two important factors might be related to the performance of computational brain models: effective dimensionality and alignment pressure. *Effective dimensionality* (ED) is a continuous measurement of the number of principal components needed to explain most of the variance in a dataset, and it is a way of estimating *latent* dimensionality in our analyses (see Figure 1a). A model with low ED encodes a relatively small number of dimensions whose variance is larger than the variance attributed to noise (i.e., whose signal-to-noise ratio (SNR) is high). In contrast, a model with high ED encodes many dimensions with high SNR. *Alignment pressure* (AP) quantifies the probability that the high SNR dimensions from a pair of subspaces will be aligned, as depicted in Figure 1b. For example, if the AP between a model subspace and the ecological subspace is high, it means that the model is likely to encode the same dimensions of image representation as those observed in the brain.

**Figure 1:**
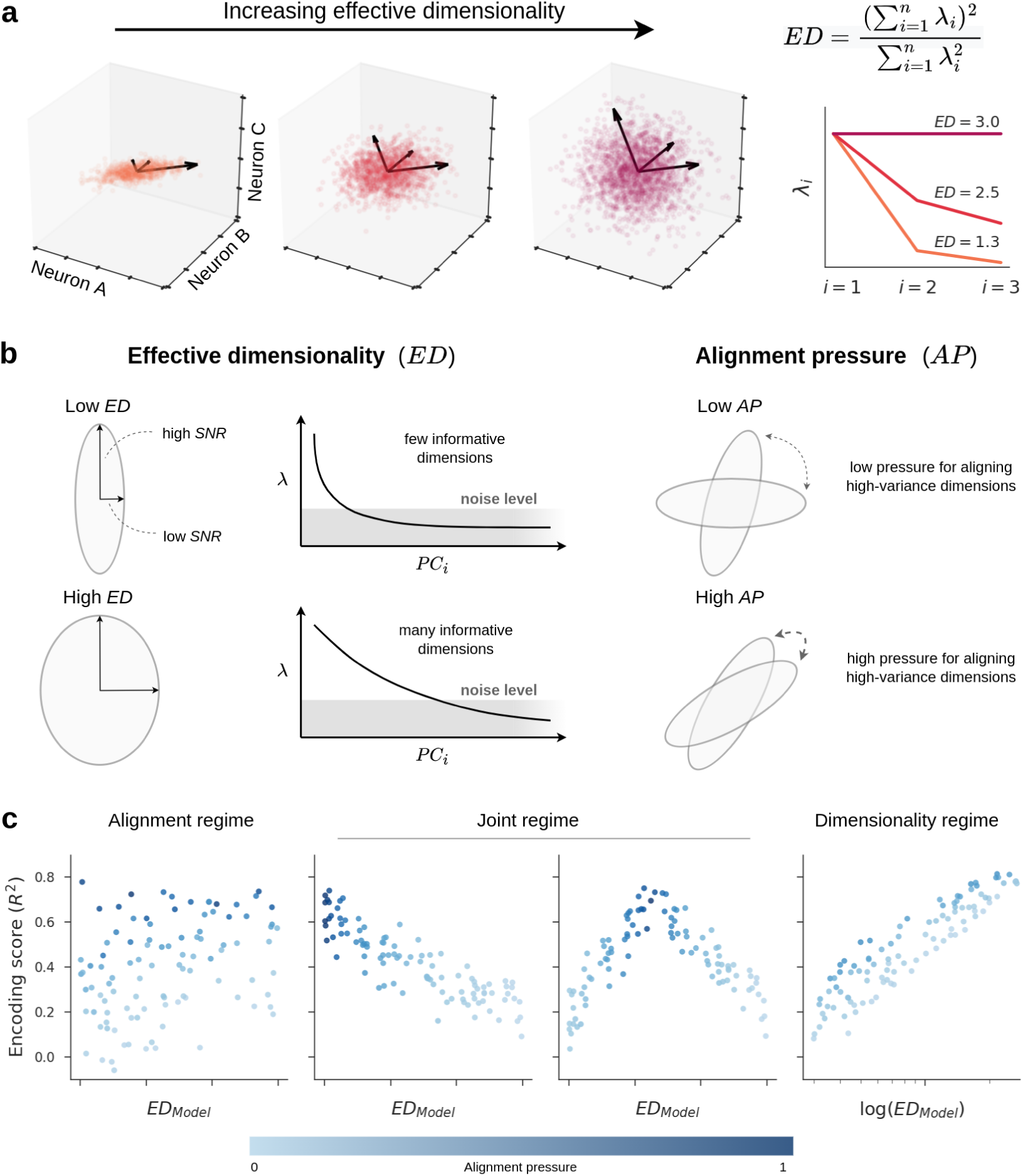
A theory of latent dimensionality and encoding performance. **a**. This panel illustrates effective dimensionality (ED) for a hypothetical population of three neurons. The data points correspond to stimuli, and the plot axes indicate the firing rates of neurons in response to these stimuli. The leftmost plot shows a scenario where the firing rates of the neurons are highly correlated and primarily extend along a single direction, resulting in an ED close to 1. The opposite scenario is shown in the rightmost plot where variance in neural responses is equally distributed across all directions, resulting in an ED of 3. On the right, we show the eigenspectra (*λ_i_*) in each scenario and the equation that describes how ED is computed from these eigenspectra. **b.** Our simulations examine two geometric properties: effective dimensionality (ED) and alignment pressure (AP). ED is a summary statistic that indicates the number of features accurately encoded by an ecological or model subspace (i.e., it is a way of estimating latent dimensionality). The eigenspectrum of a low-dimensional subspace decays quickly, suggesting that most features are dominated by noise and, therefore, poorly encoded, whereas the eigenspectrum of a high-dimensional subspace has high variance spread along a large number of dimensions. AP determines the alignment of high-variance dimensions across two subspaces. Pairs of subspaces with low AP are sampled independently with little bias for their signal dimensions to align, whereas pairs of subspaces with high AP are more likely to have substantial overlapping variance along their signal dimensions. **c.** Depending on the distributions of ED and AP in empirical models, our simulations predict different outcomes for the relationship between model ED and encoding performance. In the Alignment regime, model performance is predominantly driven by the alignment of the meaningful, signal dimensions in the model and the brain, with little to no influence of latent dimensionality. Most modeling efforts in computational neuroscience implicitly assume that models operate in the Alignment regime. Another possibility is that models operate in a Joint regime, in which there exists some optimal dimensionality at which model representations are more likely to be aligned with the brain (perhaps a low dimensionality as in the plot on the left, or a intermediate dimensionality as in the plot on the right). This is the implicit assumption behind efforts to explain brain representations with models that compress latent dimensionality (such as autoencoders). A third possibility, which has been largely overlooked, is that models operate in a Dimensionality regime, in which models with higher latent dimensionality are more likely to contain the same representational dimensions that were sampled in a neuroscience experiment. Note that the Dimensionality regime occurs when there is large variance in model ED, so we use a logarithmic scale on the x-axis for this regime.

Nearly all representational modeling efforts in computational neuroscience seek to optimize AP. For example, when researchers construct models through deep learning or by specifying computational algorithms, the hope is that the resulting model will encode representational dimensions that are strongly aligned with the representations of a targeted brain system. However, if one allows for linear transformations when mapping computational models to brain systems—a procedure that may, in fact, be necessary for evaluating such models [13]—then there are possible scenarios in which model performance can be primarily governed by ED.

To understand how ED can influence model performance, it is helpful to first consider two extreme cases. At one extreme, models with an ED of 1 can explain, at best, a single dimension of brain representation and can only do so when AP is extremely high. Such a model would need to encode a dimension that was *just right* to explain variance in a targeted brain system. At the other extreme, a model with very high ED could potentially explain many dimensions of brain representation and could do so with weaker demands on AP. This means that models with extremely high ED have a higher probability of performing well and need only be partially aligned with a targeted brain system.

The relative contributions of ED and AP will depend on their empirical distribution in actual computational models trained to predict real neural data. To better anticipate the possible outcomes, we varied our simulation parameters and identified distinct regimes for the relationship between ED, AP, and the performance of computational brain models (Figure 1c).

In the *Alignment regime*, the ED of computational models varies significantly less than their AP, such that AP predominantly drives performance in predicting neural activity. This perspective implicitly underlies most deep learning approaches for modeling visual cortex, which emphasize factors affecting the alignment between a model and the brain, such as architecture, layer depth, learning objective, and training images [e.g. 92, 13, 14, 52, 59, 93, 48, 22, 50, 97, 12, 20]. The alignment-based perspective does not entail any specific predictions about ED and, thus, suggests the null hypothesis that ED and encoding scores are unrelated (Figure 1c left panel).

Alternatively, models might inhabit a *Joint regime* where ED and AP are entangled, such that there exists some optimal dimensionality at which model representations are more likely to be aligned with the brain. Previous work has proposed that both biological and artificial vision systems gain computational benefits by representing stimuli in low-dimensional subspaces [19, 3, 58]. For instance, it has been hypothesized that dimensionality reduction along the visual hierarchy confers robustness to incidental image features [72, 23, 2, 61, 25, 24, 57]. This dimensionality-reduction hypothesis implicitly underlies a wide variety of machine learning methods that attempt to encode complex stimuli using a small set of highly informative dimensions (e.g., autoencoders) [49, 90, 96]. The strongest version of the low-dimensionality perspective predicts that ED and encoding scores will be negatively correlated or exhibit an inverted U-shape, since models with relatively low-dimensional subspaces would tend to be better aligned with the representations of visual cortex (Figure 1c middle panels).

A final possibility is that of a *Dimensionality regime*. This can occur if the computational models under consideration vary significantly in terms of ED and are sufficiently constrained to make the baseline probability of partially overlapping with visual cortex non-negligible (i.e., they have some moderate level of AP). In this case, ED will exert a strong, positive influence on expected encoding performance (Figure 1c right panel). It is unknown whether ED is a relevant factor for convolutional neural network models of visual cortex, and, if so, whether high-dimensional representations lead to better or worse models. Our simulations suggest several possible outcomes depending on the empirical distribution of ED and AP, including a previously unexplored scenario where high latent dimensionality is associated with better cross-validated models of neural activity. However, these simulations explore the effect of dimensionality in a highly idealized setting without attempting to capture the statistics of real DNNs or brains. Moreover, in our simulations, we explicitly controlled the type and level of noise in the representations, which makes the interpretation of ED straightforward, whereas, in real networks the potential contribution of noise is far more complex and there is no guarantee that ED will be related to the quality of latent dimensions. Nonetheless, these simulations can help us to generate testable hypotheses about the potential implications of latent dimensionality in real networks, which we set out to explore next. Specifically, we sought to examine the relationship between latent dimensionality and encoding performance in state-of-the-art DNNs and recordings of image-evoked responses in visual cortex.

### 2.2 Dimensionality in deep neural networks

Before presenting our analyses, it is helpful to first consider the interpretation of latent dimensionality in the context of DNNs and to highlight some important methodological details. The logic underlying dimensionality metrics like ED is that the scale of a dimension’s variance is indicative of its meaningfulness, with high variance corresponding to meaningful dimensions and low variance corresponding to random or less-useful dimensions. Even in deterministic systems, like the DNNs examined here, variance scale can indicate meaningfulness. Indeed, previous work has shown that network training expands variance along dimensions that are useful for solving tasks while leaving unaltered, or even contracting, variance along random dimensions [29, 72, 26, 41, 89]. This explains why networks trained with different random initializations end up with similar high-variance principal components but different low-variance components, and it explains why measures of network similarity appear to be more meaningful when they are weighted by variance [51, 56]. Furthermore, these empirical observations are consistent with theories of deep learning which argue that networks progressively learn the principal components of their training data and tasks, with the number of learned components increasing as a function of task and data complexity [41, 42, 36, 7, 76, 56].

It is important to note, however, that the scale of activation variance is not universally indicative of representation quality in DNNs. First, architecture-specific factors can affect ED in ways that are independent of learning, which means that ED values (and related dimensionality metrics) should only be compared across models with similar architectures [35]. Second, it is possible to arbitrarily rescale the variance of any dimension through normalization procedures (see Appendix I). Thus, to gain a better understanding of whether high-performing DNNs have expressive, high-dimensional representations or compressed, low-dimensional representations, it is important to examine the role of dimensionality while controlling for architecture. Here we controlled for architectural factors by focusing on a set of standard convolutional architectures (mostly ResNet). We provide a more detailed discussion of these points in Appendices A & I.

For our analyses, we examined a large bank of 568 layers from DNNs that varied in training task, training data, and depth. The training tasks for these DNNs included a variety of objectives, spanning both supervised (e.g., object classification) and self-supervised (e.g., contrastive learning) settings. We also included untrained DNNs. The training datasets used for these DNNs included ImageNet [75] and Taskonomy [94]. Most DNNs had ResNet50 or ResNet18 architectures [44]. We also examined a smaller set models with AlexNet [54], VGG-16 [82], and SqueezeNet [46] architectures to ensure that our findings were not idiosyncratic to ResNets. We restricted our analyses to convolutional layers at varying depths because the structure of the fully connected layers substantially differed across models. A detailed description of all models is provided in Appendix E. Using this approach, we were able to examine the effect of ED while controlling for architecture.

We empirically estimated the ED of the DNNs by obtaining layer activations in response to 10,000 natural images from the ImageNet validation set [75]. We applied PCA to these layer activations and computed ED using the eigenvalues associated with the principal components. An important methodological detail is that we applied global average pooling to the convolutional feature maps before computing their ED. The reason for this is that we were primarily interested in the variance of image *features*, which indicates the diversity of image properties that are encoded by each model, rather than the variance in those properties across space. Nevertheless, we show in Appendix F that our main results on the relationship between ED and encoding performance were observed even when ED was computed on the entire flattened feature maps without pooling (though to a lesser extent). The ED values that we computed can be interpreted as estimates of the number of meaningful dimensions of natural image representation that are encoded by each model (i.e., their latent dimensionality).

Our analyses of ED showed several general trends, which are discussed in detail in Appendix H. Briefly, we found that ED is higher for trained compared with untrained models, that ED tends to increase with layer depth, and that ED tends to be higher for models trained on a wide variety of natural images rather than only indoor scenes. These trends in ED suggest that feature expansion may be an important mechanism of the convolutional layers in DNNs.

### 2.3 Dimensionality and encoding performance for neural data

We next wanted to determine if the ED of representational subspaces in DNNs was related to their encoding performance (Figure 2). To do so, we first compared DNN models with electrophysiological recordings of image-evoked responses in macaque IT cortex—a high-level region in the ventral visual stream that supports object recognition [21]. These data were collected by Majaj et al. [62], and the stimuli in this study were images of objects in various poses overlaid on natural image backgrounds. In total, the dataset consisted of 168 multiunit recordings for 3,200 stimuli. We quantified the ability of each convolutional layer in each DNN to explain neural responses by fitting unit-wise linear encoding models using partial least squares regression, which is a standard procedure in the field for mapping computational models to neural data [93]. These encoders were evaluated through cross-validation, where regression weights are learned from a training set and then used to predict neural responses to stimuli in a held-out test set (Figure 2b). We measured encoding performance by computing the median explained variance between the predicted and actual neural responses across all recorded units.

**Figure 2:**
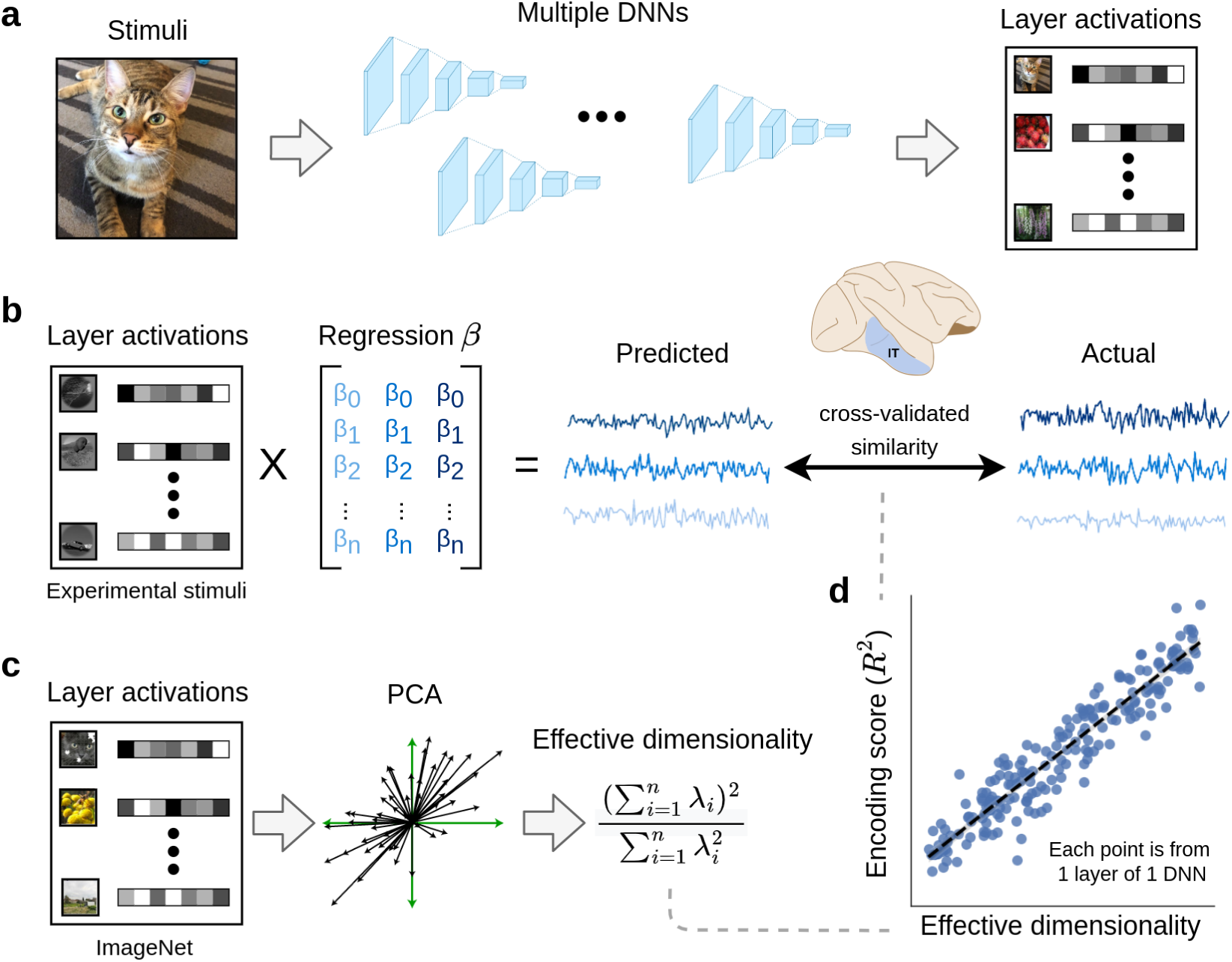
Method for comparing latent dimensionality with encoding performance for neural data. **a**. Layer activations were extracted from a large bank of DNNs trained with different tasks, datasets, and architectures. **b.** Using these layer activations as input, we fit linear encoding models to predict neural activity elicited by the same stimuli in both monkey and human visual cortex. We used cross-validation to evaluate encoding performance on unseen stimuli. **c.** To estimate the effective dimensionality of our models, we ran principal component analysis on layer activations obtained from a large dataset of naturalistic stimuli (specifically, 10,000 images from the ImageNet validation set). **d.** These analyses allowed us to examine the empirical relationship between effective dimensionality and linear encoding performance across a diverse set of DNNs and layers. DNN = deep neural network, PCA = principal component analysis.

Our analysis revealed a clear and striking effect: the encoding performance of DNN models of high-level visual cortex is strongly and positively correlated with ED (Figure 3a). This effect was also observed when separately examining subsets of models from the same DNN layer (Figure 3b), which means that the relationship between ED and encoding performance cannot be reduced to an effect of depth. (Note that the reverse is also likely true: there may be effects of depth that cannot be reduced to effects of ED.) This within-layer analysis also perfectly controls for ambient dimensionality, which is the number of neurons in a layer, and, thus, shows that this effect is specifically driven by the *latent* dimensionality of the representational subspaces. Furthermore, this effect could not be explained by categorical differences across learning paradigms or across model training datasets because it was also observed when separately examining subsets of models that were either untrained, supervised, or self-supervised (Figure 3a,b) as well as subsets of models that were trained on Taskonomy or ImageNet (Appendix F). Remarkably, this effect is not specific to the encoding model paradigm, as it was also observed when using representational similarity analysis [53], which involved no parameter fitting (see Appendix G). Finally, we also performed these analyses for human high-level visual cortex using an fMRI dataset collected by Bonner and Epstein [9]. The human fMRI results for the lateral occipital region (LO; a high-level region supporting object recognition) are shown in Figure 3c,d, and they reveal a similar positive relationship between ED encoding performance in the human brain.

**Figure 3:**
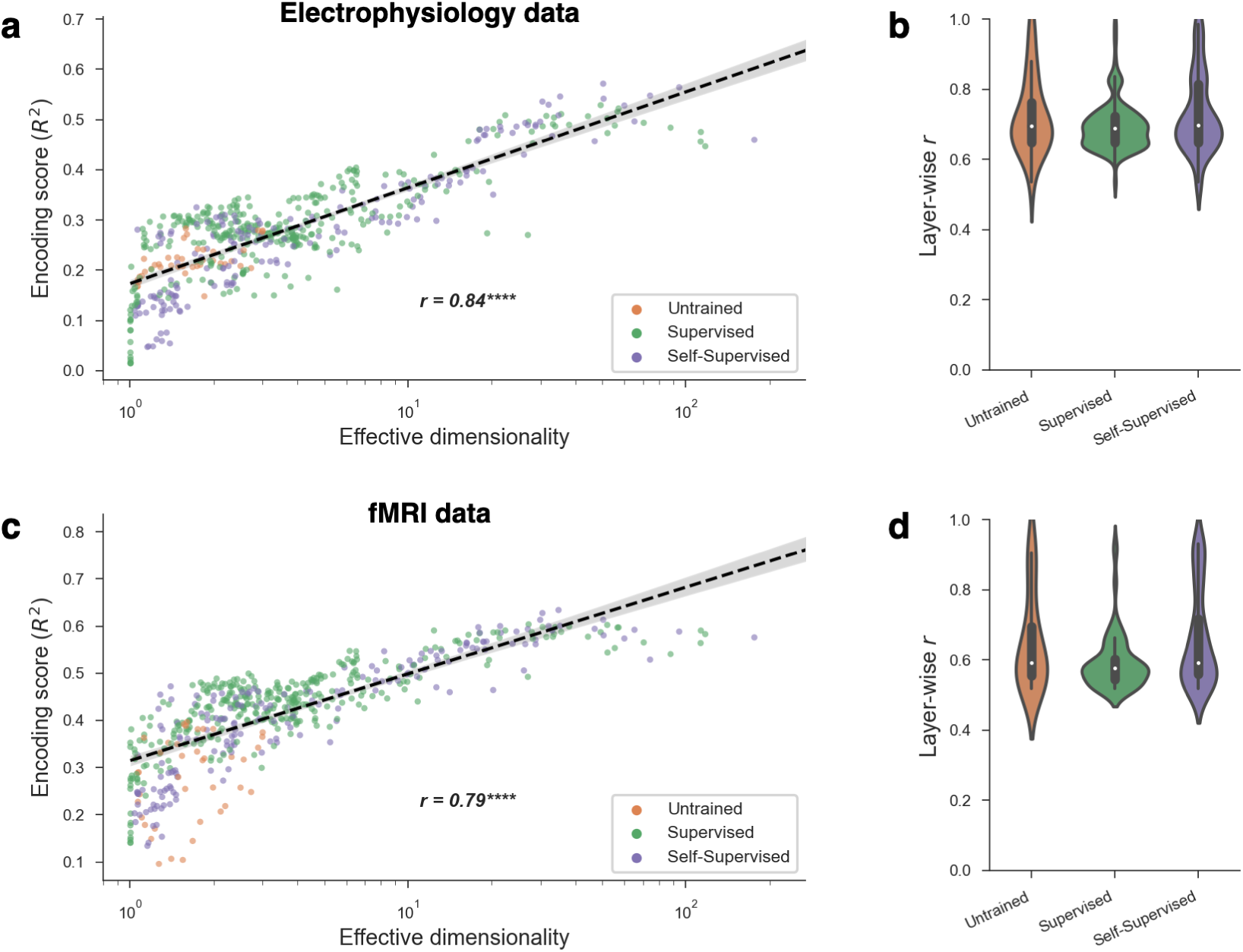
Relationship between effective dimensionality and encoding performance. **a**. The encoding performance achieved by a model scaled with its effective dimensionality. This trend also held *within* different kinds of model training paradigms (supervised, self-supervised, untrained). Each point in the plot was obtained from one layer from one DNN, resulting in a total of 568 models. Encoding performance is for the monkey IT brain region, which had the strongest relationship with ED among regions we considered. **b.** Even when conditioning on a particular DNN layer, controlling for both depth and ambient dimensionality (i.e., number of neurons), effective dimensionality and encoding performance continued to strongly correlate. The plot shows the distribution of these correlations (Pearson *r*) across all unique layers in our analyses. **c,d.** Similar results were obtained for human fMRI data.

We further performed analyses in datasets for other lower-level visual regions in monkeys (V1, V2, and V4) and for multiple other visual regions in the human brain. While the relationship between ED and encoding performance was strongest in high-level visual cortex in both monkeys and humans, similar but weaker effects were also observed in multiple other visual regions, with the exception of monkey V1 (see Appendix F. We speculate that the effect was not observed in the monkey V1 dataset for two reasons. The first is that the stimuli in this V1 dataset were simple images of synthesized textures, which may not require the complexity of high ED models Freeman et al. [28]. The second is that V1 is known to be explained by primitive edge detectors that likely emerge in most DNNs, even those with low ED. Another intriguing possibility is that each brain region could have an “optimal” value of ED where model encoding performance tends to peak (reminiscent of the “Joint regime” we showed in Figure 1c), and that this optimal value could increase along the cortical hierarchy Wang and Ponce [91]. Indeed, when we fit inverted-U shaped curves to the per-region data in Figure F.3, the results suggest that the ED with peak encoding performance increases from V1 to V2 and from V4 to IT.

In addition to ED, we examined the complete eigenspectra of all models (i.e., the variance along successive principal components). Intuitively, models with more slowly decaying eigenspectra use more of their principal components to represent stimuli. In line with this, Figure 4a shows that the more slowly a model’s eigenspectrum decays, the higher its encoding performance tends to be. Interestingly, many of the top-performing models tend to approach a power-law eigenspectrum decaying as 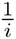, where *i* is the principal component index. This power-law decay corresponds to a proposed theoretical limit wherein representations are maximally expressive and high-dimensional while still varying smoothly as a function of changing stimuli Stringer et al. [88].

**Figure 4:**
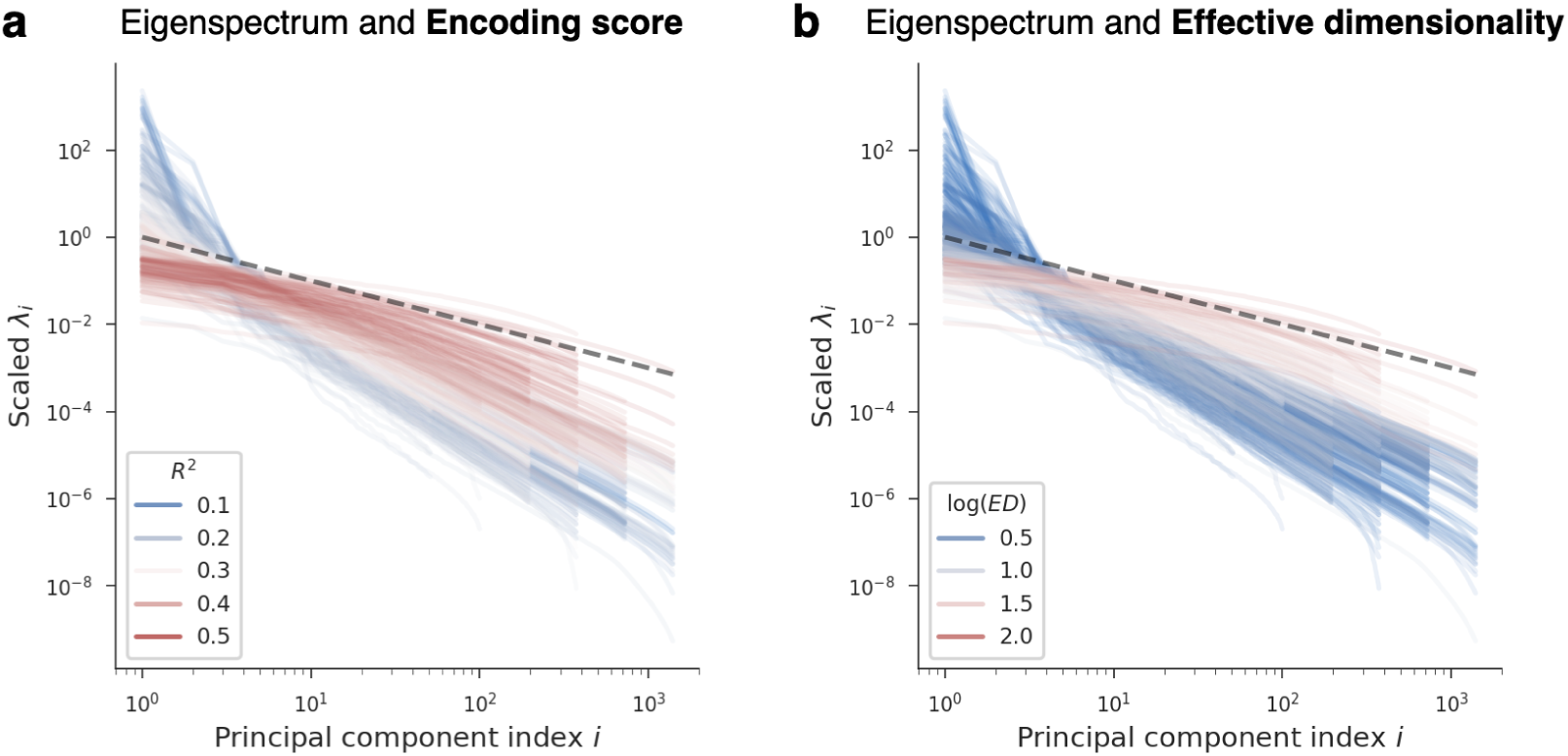
Relationship between model eigenspectra and encoding performance. Each curve shows the eigenspectrum of one layer from one DNN. The x-axis is the index of principal components, sorted in decreasing order of variance, and the y-axis is the variance along each principal component (scaled by a constant in order to align all curves for comparison). The black line is a reference for a power law function that decays as 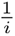, where *i* is the principal component index. This power law of 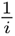 was hypothesized in Stringer et al. [88] to be a theoretical upper limit on the latent dimensionality of smooth representations. **a.** Eigenspectra are color-coded as a function of the corresponding encoding performance each model achieved. Models with more slowly decaying eigenspectra (i.e., higher latent dimensionality) are better predictors of cortical activity, with top-performing models approaching the theoretical upper bound on dimensionality proposed in Stringer et al. [88]. Encoding performance is for the IT brain region, which had the strongest relationship with ED among regions we considered. **b.** Eigenspectra are color-coded as a function of their corresponding ED. Since ED is a summary statistic of an eigenspectrum meant to quantify its rate of decay, models with more slowly decaying eigenspectra tend to have higher ED.

While visual inspection of eigenspectra plots can be illuminating, it is difficult to succinctly summarize the large amount of information that these plots contain. We, therefore, continue to use ED in our discussions below because of the concise, high-level description it provides.

Finally, in Appendix J, we compared ED with other geometric properties, and we examined how all these properties are related to both encoding performance and object classification performance. Our findings show that ED is among the strongest geometric predictors of DNN performance metrics, suggesting that high-dimensional representational subspaces allow DNNs to perform a variety of tasks related to primate vision, including the prediction of image-evoked neural responses and the classification of natural images.

In sum, these findings show that when controlling for architecture, latent dimensionality is strongly linked to the encoding performance of DNN models of high-level visual cortex, suggesting that the richness of the learned feature spaces in DNNs is central to their success as computational models of biological vision.

### 2.4 High dimensionality is associated with better generalization to novel categories

Our finding that top-performing encoding models of high-level visual cortex tend to have high dimensionality was surprising given that previous work has either not considered latent dimensionality [92, 13, 14, 52, 59, 93, 48, 22, 50, 97, 12, 20] or argued for the opposite of what we discovered: namely that low-dimensional representations better account for biological vision and exhibit computational benefits in terms of robustness and categorization performance [3, 58]. We wondered whether there might be some important computational benefits of high-dimensional subspaces that have been largely missed in the previous literature. Recent theoretical work on the geometry of high-dimensional representations suggests some hypotheses [85, 88, 55]. Specifically, it has been proposed that increased latent dimensionality can improve the learning of novel categories, allowing a system to efficiently generalize to new categories using fewer examples [85]. Efficient learning is critical for visual systems that need to operate in a diversity of settings with stimulus categories that cannot be fully known *a priori*, and it is something that humans and other animals are remarkably good at [71, 83, 6, 15].

Thus, we examined whether the dimensionality of our DNNs was related to their generalization performance on newly learned categories. We employed a standard transfer learning paradigm in which we fixed the representations of our DNN models and tested whether they could generalize to a new target task using only a simple downstream classifier (depicted in Figure 5a). Following the approach in Sorscher et al. [85], we trained a classifier on top of each model’s representations using a simple prototype learning rule in which stimuli were predicted to belong to the nearest class centroid. We used 50 object categories from the ImageNet-21k dataset [74] and trained on 50 images per category, evaluating on another 50 held-out images from the same categories. Importantly, none of the ImageNet-21k classes appeared in any of the datasets our models were pre-trained on, allowing us to assess a relationship between the latent dimensionality of a representation and its ability to classify novel categories. The results, illustrated in Figure 5b, show a striking benefit of high-dimensionality for this task. Even though high-dimensional representations have traditionally been thought to be undesirable for object classification [72, 23, 25, 24, 57, 17], they proved to be extremely effective in separating novel categories. This suggests that while low-dimensional representations may be optimal for performing specialized tasks (such as separating the fixed set of categories in the standard ImageNet training set), high-dimensional representations may be more flexible and better suited to support open-ended tasks [45, 30, 85, 26, 11].

**Figure 5:**
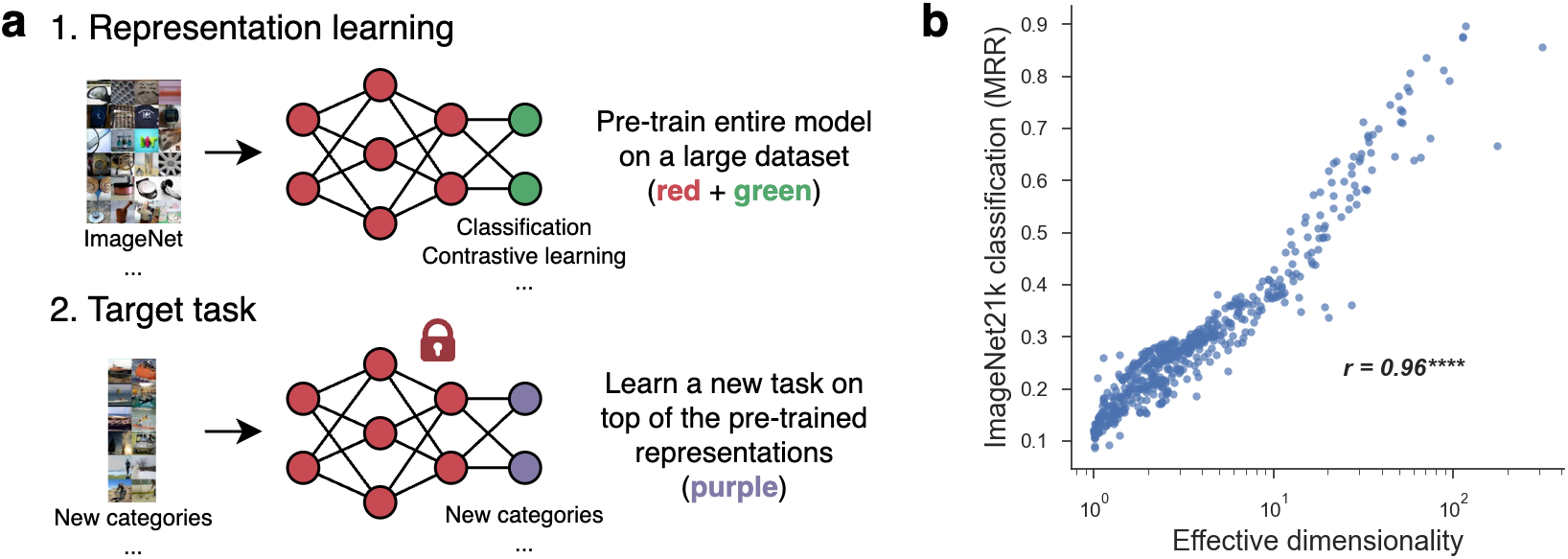
The computational benefit of high effective dimensionality in generalization to new object categories. We examined the hypothesis that high-dimensional representations are better at learning to classify new object categories [85]. **a.** We tested this theory using a transfer learning paradigm, where our pre-trained model representations were fixed and used to classify novel categories through a prototype learning rule. **b.** High-dimensional models achieved substantially better accuracy on this transfer task, as measured using the mean reciprocal rank (MRR).

### 2.5 High dimensionality concentrates projection distances along linear readout dimensions

How do high-dimensional models achieve better classification performance for novel categories? If we consider an idealized scenario in which category instances are distributed uniformly within unit spheres, it is a geometric fact that projections of these subspaces onto linear readout dimensions will concentrate more around their subspace centroids as dimensionality increases [85, 39, 40]. The reason for this is that in high dimensions, most of the subspace’s mass is concentrated along its equator, orthogonal to the linear readout dimension. This blessing of dimensionality is typically referred to as the *concentration of measure phenomenon* [39], and we depict it for the case of an idealized spherical subspace in Figure 6a,b (see Section 4, Materials and Methods).

**Figure 6:**
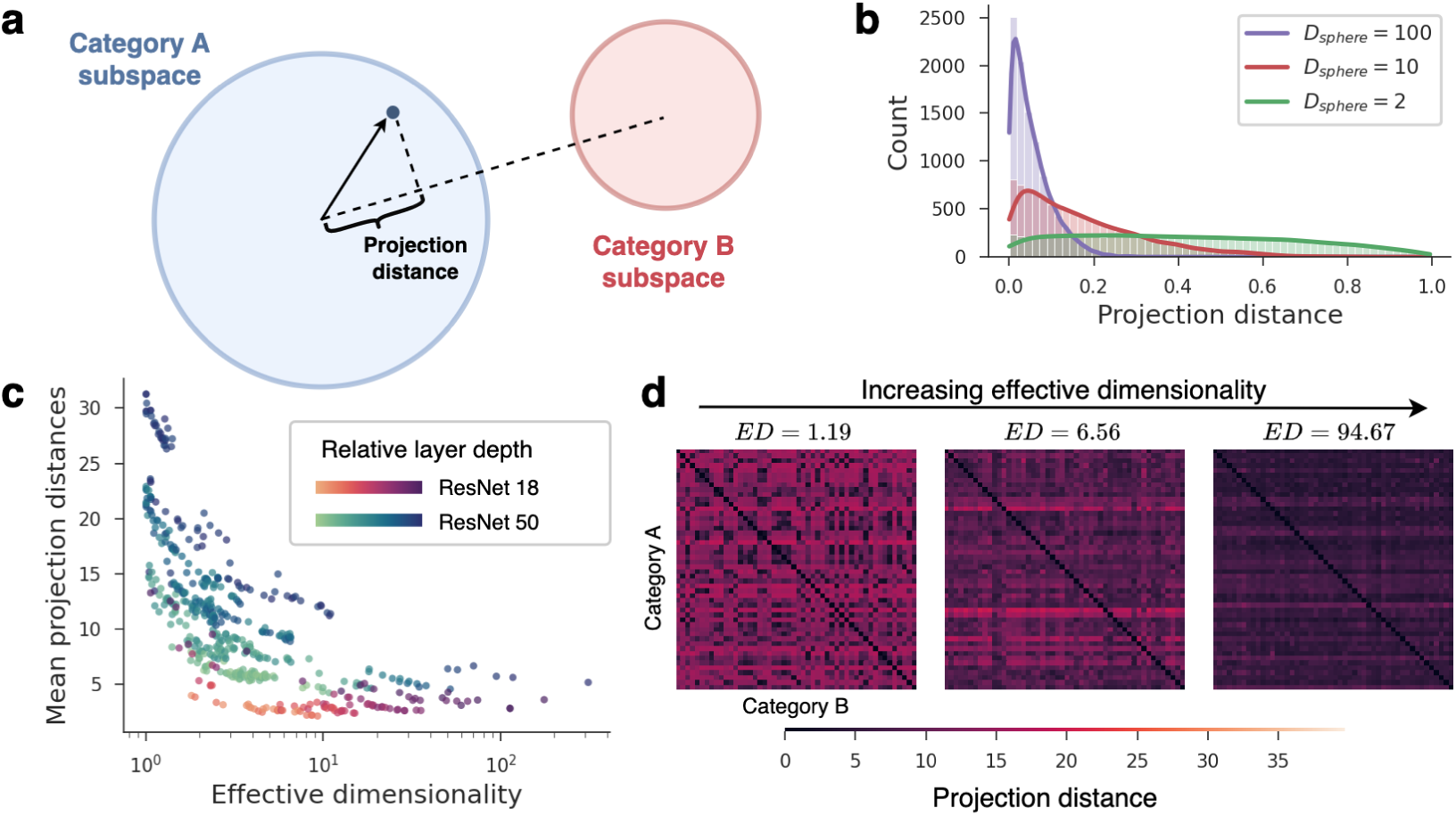
High-dimensional models concentrate sample projections close to their class centroids. **a**. For binary classification, the projection distance of a sample refers to the sample’s distance from its class centroid along the classification readout direction, normalized by the subspace radius. **b.** For idealized spherical subspaces, the distribution of projection distances concentrates more tightly around 0 as the dimensionality *D_sphere_* increases. **c.** Empirically, the mean projection distances of our models decreased as effective dimensionality increased, matching what is predicted by theory. Note that because the magnitude of projections partially depend on the model architecture and layer depth (denoted by different colors), projection distances form distinct bands in the plot. However, when looking only at models with the same architecture and layer (i.e., looking at points sharing the same color), projection distances reliably decrease with ED. **d.** Full projection distance matrices, computed along classification readout directions between all object category pairs. Matrices are shown for three different models of increasing effective dimensionality.

Although we do not know the geometric shapes of category subspaces in DNNs or whether they can be approximated by elliptical subspaces, we can, nonetheless, test whether there is empirical evidence that a similar concentration phenomenon occurs in our models. To answer this question, we computed the average sample projection distance between every pair of our 50 ImageNet-21k classes, normalized by an estimate of the subspace radius for each class (see Section 4, Materials and Methods). This yielded a matrix of all pairwise projection distances for each model. Figure 6c shows that, as predicted, the mean projection distance systematically decreases as model ED increases. This means that sample projection distances concentrate closer to class centroids as model dimensionality increases, allowing DNNs with higher ED to discriminate novel categories more effectively. This concentration effect exemplifies an underappreciated computational advantage conferred by the geometric properties of high-dimensional subspaces.

## 3 Discussion

By computing geometric descriptors of DNNs and performing large-scale model comparisons, we discovered a geometric phenomenon that has been overlooked in previous work: DNN models of high-level visual cortex benefit from high-dimensional latent representations. This finding runs counter to the view that both DNNs and neural systems benefit by compressing representations down to low-dimensional subspaces [3, 72, 23, 25, 24, 57, 70, 96, 90, 38, 2, 61, 49, 18, 33, 67, 64, 58, 31, 77, 34].

Furthermore, our findings suggest that DNNs models of high-level visual cortex are best understood in terms of the richness of their natural image representations rather than the details of their training tasks and training data.

Our results speak to a fundamental question about the dimensionality of neural population codes. Empirical estimates have consistently shown that the latent dimensionality of both DNNs and neural systems is orders of magnitude lower than their ambient dimensionality (i.e., the number of neurons they contain) [23, 19, 3, 38, 18, 33, 34, 67, 64, 58, 32]. Furthermore, there are compelling theoretical arguments for the computational benefits of low-dimensional codes, which may promote robustness to noise [72, 96, 57, 24, 2, 61], abstraction and invariance [49, 90, 25, 31, 32], compactness [19], and learnability for downstream readouts following earlier network layers such as V1 [1, i.e., avoiding the curse of dimensionality]. It is, thus, no surprise that many neuroscientists and machine learning researchers have argued that a signature of optimal population codes is the degree to which they reduce latent dimensionality, focusing on the minimum components needed to carry out their functions. However, there are competing theoretical arguments for the benefits of high-dimensional population codes. High-dimensional codes are more efficient [66, 4], they allow for the expression of a wider variety of downstream readouts [30], and they may have counterintuitive benefits for learning new categories due to concentration phenomena in high dimensions [85]. Furthermore, recent evidence from large-scale data in mouse visual cortex suggests that cortical population codes are higher dimensional than previously reported and may, in fact, approach a theoretical limit, above which the code would no longer be smooth [88]. Our findings provide strong evidence for the benefits of high-dimensional population codes. Specifically, we demonstrated two major benefits that are directly relevant to computational neuroscientists. First, high-dimensional DNNs provide more accurate cross-validated predictions of cortical image representations. In fact, when looking at the eigenspectra of our top-performing models, they appear to approach the upper limit on dimensionality that was proposed in Stringer et al. [88]. Second, high-dimensional DNNs are more effective at learning to classify new object categories. We suggest that while low-dimensional codes may be optimal for solving specific, constrained tasks on a limited set of categories, high-dimensional codes may function as general-purpose representations, allowing them to better support an open-ended set of downstream tasks and stimuli.

Our findings also have implications for how neuroscientists interpret the relationship between DNNs and neural representations. When developing theories to explain why some DNNs are better than others as computational brain models, it is common for researchers to place a strong emphasis on the task on which the network was optimized [92, 13, 14, 52, 59, 93, 48, 22, 50, 97, 12, 20]. We and others have found that a variety of tasks are sufficient to yield high-performing encoding models of visual cortex [22, 50, 97, 20]. However, when analyzing the geometry of these networks, we found that a common thread running through the best-performing models was their strong tendency to encode high-dimensional subspaces. It is worth emphasizing that we were only able to observe this phenomenon by analyzing the geometry of latent representations in many DNNs with the same architecture and examining large-scale trends across these models. In other words, the most important factors for explaining the performance of these models were not evident in their task-optimization constraints but rather in their latent statistics. This finding is consistent with other recent work showing that widely varying learning objectives and architectures—including transformer architectures from computational linguistics—are sufficient to produce state-of-the-art encoding performance in visual cortex, which suggests that task and architecture are not the primary explanation for the success of DNNs in visual neuroscience [50, 97, 20]. Our findings are also consistent with recent work that calls into question the apparent hierarchical correspondence between DNNs and visual cortex [80, 86]. Indeed, we found that the relationship between latent dimensionality and encoding performance generalized across layer depth, meaning that even within a single layer of a DNN hierarchy, encoding performance can widely vary as a function of latent dimensionality. Our work suggests that the geometry of latent representations offers a promising level of explanation for the performance of computational brain models.

A recent preprint examined the ED of layer activations across a highly varied set of neural networks and found that while ED was higher for trained versus untrained versions of the same architectures, as was also shown here, variation in ED across diverse trained networks was not correlated with encoding performance in visual cortex [20]. However, we want to emphasize that this latter finding is not unexpected given the nature of the model set examined in the study. Specifically, the authors examined the best performing layer from models with diverse architectures and varied normalization layers, including convolutional networks, transformers, and multilayer perceptrons, all of which can have architecture-specific effects on the shape of representational eigenspectra (see Results 2.2 and Appendix I)). Importantly, the existence of such architecture-specific effects means that a given ED value will not have the same meaning across all networks, and, for this reason, we caution against comparing ED values (and related dimensionality metrics) across networks with highly varied architectures [35]. Furthermore, the ED metric computed in Conwell at al. is different from the ED metric that we found to be most informative in our own analyses. Specifically, we computed ED based on channel covariance by using global pooling, whereas Conwell at al. computed ED based on a mixture of both channel and spatial covariance, which in our models predicts encoding performance less well (Appendix F). We focused on channel covariance because it reflects the richness of feature types in a network and because it has been shown to be highly informative for theories of learning in convolutional architectures [41]. In contrast, spatial covariance reflects the spatial smoothness of layer activations, which is informative in its own right but appears to be less useful for understanding the richness of DNN representations. In sum, we argue that the most informative approach for understanding the role of dimensionality in DNNs is to examine how variations in dimensionality are related to variations in model performance while controlling for architecture. We further propose that channel covariance, rather than spatial covariance, offers the best lens on the key representational properties of DNNs.

We should also note that because we focused specifically on channel covariance in our main analyses, our notion of dimensionality is different from the dimensionality metrics used in most previous studies of CNNs, which generally did not make a distinction between channel and spatial information [e.g. 3, 19]. We believe that channel and spatial covariance have different functional consequences for visual representations, and it may be important to keep the distinction between these two factors in mind when examining dimensionality and when comparing results across studies. Nonetheless, we did not find evidence in our study to suggest that dimensionality trends in CNNs are markedly different when spatial information is included—-rather when including spatial information in our analyses, we found qualitatively similar, though weaker, relationships between dimensionality and measures of model performance. We suggest that a useful direction for future work is to better understand the specific ways in which channel and spatial covariance might differentially affect task performance in neural networks.

Our results raise several important considerations for future work. First, while our findings show that computational models of visual cortex benefit from high latent dimensionality, our method cannot speak to the dimensionality of visual cortex itself and was not designed to do so. Indeed, the theory that we presented in Section 2.1 predicts that high-dimensional DNNs should generally better explain neural activity *even if neural representations in the brain are low-dimensional*. However, our findings suggest a promising model-guided approach for tackling this issue: one could use high-dimensional DNNs to create stimulus sets that vary along as many orthogonal dimensions as possible. This sets up a critical test of whether the latent dimensionality of visual cortex scales up to the dimensionality of the model or, instead, hits a lower-dimensional ceiling.

Second, we found that one way in which DNNs can achieve both strong encoding performance and strong image classification performance is by increasing the latent dimensionality of their representations. However, this finding diverges from previous work that has linked better classification performance to dimensionality reduction in DNN representations [3, 72]. We believe that this discrepancy arises due to a fundamental problem with classification metrics: DNNs with the best classification scores are optimal for a single task on a small and closed set of categories (e.g., ImageNet classes), but these seemingly optimal DNNs may be less useful for representing new categories or for representing meaningful variance within a category (e.g., object pose). This problem with classification metrics may help to explain why the strong correlation between DNN classification performance and cortical encoding performance [93, 97] appears to break down at the highest levels of classification accuracy [78, 79, 60] (see an extended discussion of these issues in Appendix A). Future work should move beyond classification accuracy and explore richer descriptions of representational quality.

Third, it is important to keep in mind that ED can vary as a function of architecture-specific factors and that some classes of operations can break the relationship between ED and the diversity of learned features, as we show in Appendix I. Thus, high ED is not universally associated with better DNN performance and comparisons of ED across different architectures may not be informative. In our analyses, we were able to reveal the benefits of high latent dimensionality by controlling for architecture. An exciting direction for future work is the development of new metrics of representational richness whose interpretation is independent of architecture.

Finally, an open question is whether our results are specific to convolutional neural networks and higher visual cortex or whether similar results could be obtained for other classes of computational models (e.g., transformers) and other sensory and cognitive domains (e.g., audition, language). Note that our approach of using global pooling to focus on channel covariance means that our methods are tailored to convolutional architectures. Thus, further work will be needed to apply our approach to other classes of architectures in which channel and spatial information are not readily separable.

In sum, we propose that the computational problems of real-world vision demand high-dimensional representations that sacrifice the competing benefits of robust, low-dimensional codes. In line with this prediction, our findings reveal striking benefits for high dimensionality: both cortical encoding performance and novel category learning scale with the latent dimensionality of a network’s natural image representations. We predict that these benefits extend further and that high-dimensional representations may be essential for handling the open-ended set of tasks that emerge over the course of an agent’s lifetime [45, 30, 85, 26, 11].

## 4 Materials and Methods

### Simulations

The theory and rationale behind our simulations is explained in Appendix B. Precise implementation details are provided in Appendix C.

### Deep neural networks

We used 46 different DNNs, each with either a ResNet18, ResNet50, AlexNet, VGG-16, or SqueezeNet architecture. Training tasks included supervised (e.g., object classification) and self-supervised (e.g., colorization) settings. We also used untrained models with randomly initialized weights. The training datasets of these DNNs included ImageNet [75] and Taskonomy [94]. Further details describing each model are provided in Appendix E. Convolutional layers in ResNets are arranged into 4 successive groups, each with a certain number of repeated computational units called blocks. We extracted activations from the outputs of each of these computational blocks, of which there are 8 in ResNet18 and 16 in ResNet50. For other architectures, we used layers that were close to evenly spaced across the depth of the model. Across our 46 DNNs, this resulted in a total of 568 convolutional layers that we used for all further analyses.

### Neural datasets

Neural responses were obtained from a publicly available dataset collected by Majaj et al. [62]. Two fixating macaques were implanted with two arrays of electrodes in IT—a visual cortical region in later stages of the ventral-temporal stream—resulting in a total of 168 multiunit recordings. Stimuli consisted of artificially-generated gray-scale images composed from 64 cropped objects belonging to 8 categories, which were pasted atop natural scenes at various locations, orientations, and scales. In total, the dataset held responses to 3,200 unique stimuli.

In Appendix F, we also show results on additional datasets. The V4 electrophysiology dataset was collected in the same study as for IT [62]. The V1 electrophysiology dataset was collected by Freeman et al. [28], and consisted of responses to 9000 simple synthetic texture stimuli. In addition to our electrophysiology datasets, we also used a human fMRI dataset collected by Bonner and Epstein [9]. The stimulus set consisted of 810 objects from 81 different categories (10 object tokens per category). fMRI responses were measured while 4 subjects viewed these objects, shown alone on meaningless textured backgrounds, and performed a simple perceptual task of responding by button press whenever they saw a “warped” object. Warped objects were created through diffeomorphic warping of object stimuli [87]. The methods for identifying regions of interest in these data are detailed in Bonner and Epstein [9]. The localizer scans for these data did not contain body images, and, thus, a contrast of faces-vs.-objects was used to select voxels from the parcel for the extrastriate body area (EBA).

### Predicting neural responses

We obtained activations at a particular layer of a DNN to the same stimuli that were used for obtaining neural responses. The output for each stimulus was a three-dimensional feature map of activations with shape *channels × height × width*, which we flattened into a vector. For our monkey electrophysiology dataset, we fit a linear encoding model to predict actual neural responses from the DNN layer features through partial-least-squares regression with 25 latent components, as in Yamins et al. [93] and Schrimpf et al. [78]. To measure the performance of these encoding models, we computed the Pearson correlation between the predicted and actual neural responses on held-out data using 10-fold cross validation, and averaged these correlations across folds. We aggregated the per-neuron correlations into a single value by taking the median, which we then normalized by the median noise ceiling (split-half reliability) across all neurons. This normalization was done by taking the squared quotient *r*^2^ = (*r/r_ceil_*)^2^, converting our final encoding score into a coefficient of explained variance relative to the noise ceiling. The entire process described above for fitting linear encoding models was implemented with the Brain-Score package [78, 79] using default arguments for the Majaj et al. [62] public benchmark.

The process for fitting voxel-wise encoding models of human fMRI data (presented in Appendix F) differed slightly from the above. For each of our 4 subjects, we used 9-fold cross-validated ordinary least squares regression. Encoding performance was measured by taking the mean Pearson correlation between predicted and held-out voxel responses across folds, and then aggregated by taking the median across voxels. Finally, this score was averaged across subjects. No noise-ceiling normalization was used.

Before fitting these linear encoding models, we also applied PCA to the input features in order to keep the number of parameters in the encoders constant. For each model layer, principal components were estimated using 1000 images from the ImageNet validation set. Layer activations to stimuli from the neural dataset were then projected along 1000 components and finally used as regressors when fitting linear encoding models. We emphasize that this dimensionality reduction procedure was done purely for computational reasons, as using fewer regressors reduced the computer memory and time required to fit our encoding models. Our findings are not sensitive to this particular decision, as we obtained similar results by applying average-pooling instead of PCA to our DNN feature maps as an alternative method for reducing the number of regressors.

### Estimating latent dimensionality

We used a simple linear metric called effective dimensionality (ED) [10, 27, 69, 37] to estimate the latent dimensionality of our model representations. ED is given by a formula that quantifies roughly how many principal components contribute to the total variance of a representation. We, thus, ran PCA on the activations of a given model layer in response to a large number of natural images (10,000 from the ImageNet validation set) in an effort to accurately estimate its principal components and the variance they explain. An important methodological detail is that we applied global average-pooling to the convolutional feature maps before computing their ED. The reason for this is that we were primarily interested in the variance of image *features*, which indicates the diversity of image properties that are encoded by each model, rather than the variance in those properties across space.

### Classifying novel object categories

To see how model ED affected generalization to the task of classifying novel object categories, we used a transfer learning paradigm following Sorscher et al. [85]. For a given model layer, we obtained activations to images from *M* = 50 different categories each with *N_train_* = 50 samples. We then computed *M* category prototypes by taking the average activation pattern within each category. These prototypes were used to classify *N_test_* = 50 novel stimuli from each category according to a simple rule, in which stimuli were predicted to belong to the nearest prototype as measured by Euclidean distance. This process was repeated for 10 iterations of Monte Carlo cross-validation, after which we computed the average test accuracy. Importantly, none of these object categories or their stimuli appeared in any of the models’ pre-training datasets. Stimuli were taken from the ImageNet-21k dataset [74], and our object categories were a subset of those used in Sorscher et al. [85].

### Projection distances along readout dimensions

Section 2.5 investigated how points sampled from high-dimensional object subspaces project along linear readout vectors. First, we illustrated this phenomenon in Figure 6b using simulated data. We sampled *N* points uniformly in the volume of a sphere of dimensionality *d* and radius 1. For each point, we projected it along a random unit vector in R*^d^*, forming a distribution of projection distances from samples in the sphere to its centroid. We then plot this distribution for increasing values of *d*.

For the experimental results in Figures 6c,d, we sampled 50 random images from the same 50 ImageNet-21k object categories described earlier. For every pair of object categories *i* and *j*, we created a classification readout vector **w***^i,j^* by taking the difference between category centroids normalized to unit length:

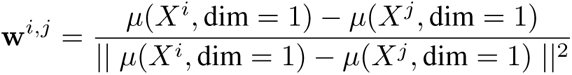

where 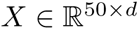 is a matrix of model activations for a particular object category and 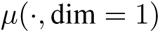 computes the category centroid by taking the mean along the row dimension. This is the readout vector along which samples would be projected using a prototype learning classification rule. We then projected each sample *k* in category *i* along the readout vector, yielding a scalar projection distance 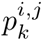 to the category centroid:

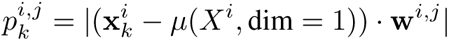

For each pair of categories (*i, j*), we therefore had a vector of projection distances 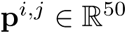 coming from the 50 image samples in category *i*, and we took the average of this vector to give a mean projection distance. This mean projection distance was the summary statistic we were interested in, since theory predicts that it is both influenced by dimensionality and relevant to classification [85]. However, one issue is that we wished to compare this summary statistic across different architectures and layers, which might have different feature scales. To correct for this, we needed to normalize each model’s mean projection distances by some normalizing factor that quantifies the feature scale. We computed this normalizing factor by taking the square root of the average variance for each category representations, which conceptually can be thought of as quantifying category subspace’s “radius” [85]. Specifically, for a given model, we computed:

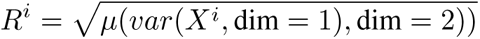

where 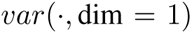 computes the per-dimension variance. We refer to the resulting values as the mean normalized projection distances 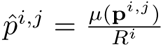. For *i*, *j* =1,…50 object categories, this procedure yields a 50 *×* 50 matrix 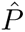 for each model. Figure 6d shows examples of these matrices for models of increasing ED. The means of these matrices as a function of ED are shown for all models in Figure 6c.

## Data availability

The neural electrophysiology datasets are publicly available and was collected in prior work by Majaj et al. [62] and Freeman et al. [28]. It is imported automatically using the BrainScore version 0.2 Python library through code in our publicly available project repository. The fMRI data was collected in prior work by Bonner and Epstein [9] and is publicly available here: https://osf.io/ug5zd/.

## Code availability

All code used for this project has been made publicly available on GitHub https://github.com/EricElmoznino/encoder_dimensionality.

## Acknowledgments and Disclosure of Funding

The brain icon in Figure 2 was obtained from Smith Breault [84].

## Appendix A Anticipated questions

In this section, we address some questions that readers are likely to have regarding our results and conclusions. While certain parts of our responses are more speculative in nature and have yet to be tested empirically, we believe that they nevertheless provide useful insights.

### Question: Isn’t it trivially true that models with higher latent dimensionality will exhibit better encoding performance?

It is important to emphasize that the relationship between ED and encoding performance cannot be explained as a trivial statistical consequence of models with high ED. First, all models were evaluated using cross-validation, which means that the only way for a model to perform well is by explaining meaningful variance that generalizes to held-out data. If models with high ED were simply overfit to the training data, their performance on the held-out test data would be poor. Second, our ED metric characterizes the distribution of variance in the eigenspectrum, but it does not directly indicate the number of available dimensions in a system, nor does it change the number of parameters in the model. In fact, all models examined here were full rank, meaning that their image representations spanned the maximum number of latent dimensions. Thus, in our analyses, ED alone has no direct relationship to the maximum number of latent dimensions that could potentially be used in a regression. Finally, the data that we modeled come from a high-level visual region (IT) whose image-evoked responses have long been a challenging target for computational modelers. In fact, decades of efforts to model the representations in this brain region directly led to the advent of deep learning approaches for the computational neuroscience of vision [93, 52]. If any model with high ED could trivially explain the representations of IT, then neuroscientists would have no need for deep neural networks. One could, instead, solve the challenging problem of modeling IT by running linear regression on RGB pixel values and adding polynomials or interaction terms until ED was high enough to account for the variance in neural responses. The reason that such an approach would not work is that the space of all possible image representations is infinite: there is an unlimited variety of arbitrary computations that could be used to add dimensions to a model. Models that achieve high ED through arbitrary computations would have a negligible probability of overlapping with the representations of visual cortex. We, thus, suspect that the use of performance-optimized DNN architectures is critical for constraining the computations of encoding models and increasing their overlap with cortical representations.

### Question: Why should we care about latent dimensionality if ImageNet classification performance also correlates with encoding performance?

Prior work has found that a model’s object classification accuracy (e.g., on ImageNet) strongly correlates with its encoding performance for certain brain regions [93, 97]. Could we simply focus on classification accuracy, or is latent dimensionality theoretically important in its own right?

While object classification accuracy seems to account for the encoding performance of current models, it is worth asking whether or not it can be a viable theory going forward. To say that it is a complete explanation is to say that we believe the correlation between classification accuracy and encoding performance will hold indefinitely, across the space of all current and future models (i.e., a model that performs better on object classification will always obtain higher encoding performance). This is unlikely to be the case. Optimal object classification requires that a model be invariant to features unrelated to object identity, such as orientation and position, which can only contribute to noise in the classifier [72]. However, we know that the brain represents orientation, position, and a host of other features unrelated to object identity. Therefore, we know that the object classification theory of encoding performance breaks down in some regime, and that the true dimensionality of visual cortex must be higher than what ideal object classification models would predict. Indeed, initial results suggest that the relationship between object classification and encoding performance does indeed break down past a certain ImageNet classification accuracy [78, 79]. A theory based on latent dimensionality (and alignment pressure) has the potential to explain the encoding performance of both current and future models on more rich neural datasets, and it may help us to understand why the relationship between encoding performance and classification accuracy breaks down at the highest levels of classification performance.

An interesting question emanating from this discussion is whether the observed relationship between classification accuracy and encoding performance might be overly optimistic due to the limited space of DNNs that are available for computational neuroscientists to examine. Most of the DNNs in visual neuroscience are trained on ImageNet or similar image databases, and we do not have DNNs that can perform open-ended tasks in complex, real-world environments. If we did have DNNs that handled more complex and naturalistic visual behaviors, we postulate that they would surpass the encoding performance of our best object-classification models (and also have higher dimensionality). With the current space of state-of-the-art DNNs being dominated by (a) supervised object classification and (b) self-supervised objectives that learn invariances tailored to object classification, we are bound to observe the current correlation between object classification performance and encoding performance because object recognition is undoubtedly *one* important problem that biological vision solves—but, importantly, it is one of *many* complex problems solved by the representations of visual cortex.

Finally, ED and classification performance are two fundamentally different *levels* of explanation that are mutually compatible because high classification accuracy itself is explained by certain geometric properties of a representation, of which ED is an important one (see Sections 2.4 and 2.5). We elaborate on this point in Appendix J where we examine the relationship between classification performance, encoding performance, and ED in our models.

### Question: Does effective dimensionality really represent the number of accurately encoded visual features?

Essential to our theory is the assumption that the variance of a representation along a particular dimension is proportional to the meaningfulness of the feature it encodes. In such cases, it is valid to say that ED roughly quantifies the number of encoded visual features. This interpretation is central to popular dimensionality reduction techniques, such as PCA, and it has good theoretical support given that high-variance dimensions are typically more robust and would, thus, be best-suited for carrying the signal in a population code. Importantly, in the DNN literature, recent findings have shown that neural networks expand variance along dimensions that are useful for solving their tasks and contract variance along noise dimensions that are left over from their random initialization [29, 72, 26]. There is, thus, a straightforward relationship between the number of meaningful latent dimensions in a neural network and the shape of the principal component variance spectrum.

Nonetheless, there is no guarantee that high-variance dimensions correspond to meaningful signal and that low-variance dimensions correspond to random features or noise. Furthermore, ED is only an imperfect summary statistic of the rate of decay of the eigenspectrum, and does not directly quantify the number of meaningful features. An interesting direction for future work would be the development of dimensionality metrics that try to explicitly differentiate between meaningful and random dimensions in DNNs, as can be done for neural data when using repeated stimulus presentations [88].

## Appendix B Theory of latent dimensionality and encoding performance

While the space of all possible visual stimuli is vast and high-dimensional, we can define many lower-dimensional subspaces within it, referred to as *subspaces* B.1a. For instance, the *natural image subspace* consists solely of images taken from the physical world, and typical neuroscience experiments consist of tightly controlled stimulus sets spanning a *data subspace*. In a similar way, we can formalize visual representations and the features they encode using the framework of subspaces embedded in a higher-dimensional visual space.

**Figure B.1:**
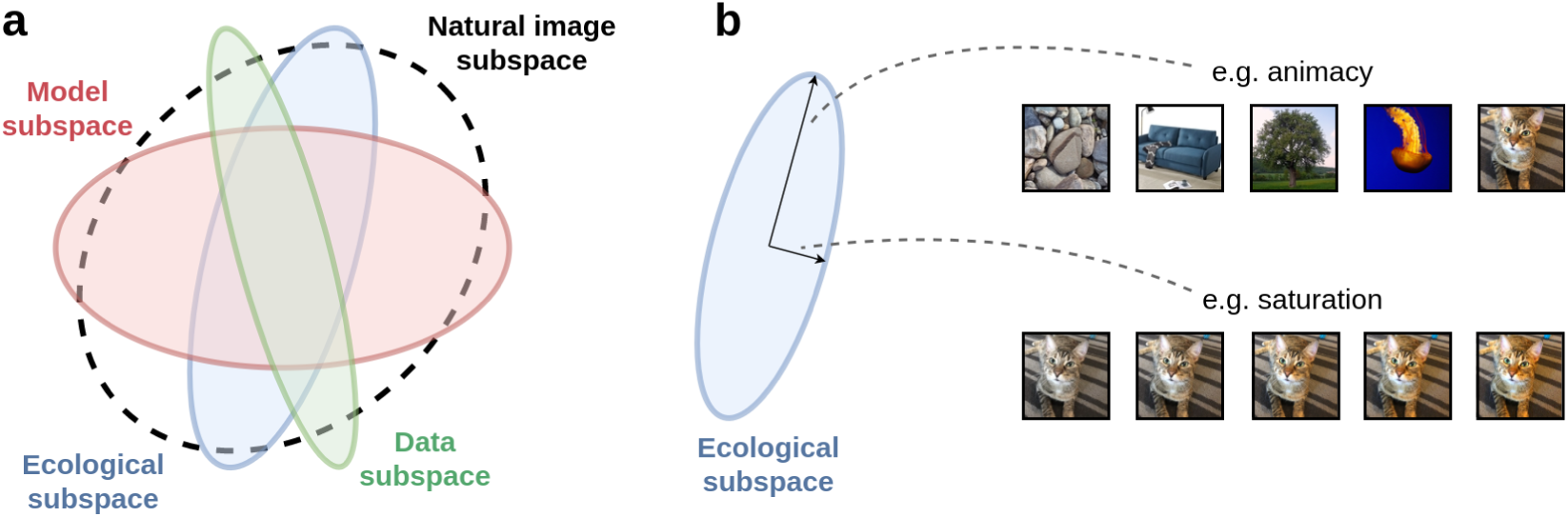
A theory of latent dimensionality and encoding performance. **a**. Our theory models the distribution of natural images, the distribution of experimental stimuli, and the features encoded by brains and models as lower-dimensional subspaces embedded in a high-dimensional ambient space (denoted here using ellipses of varying eccentricity). **b.** For ecological and model subspaces, the variance along a dimension represents the accuracy with which it is encoded. For example, visual cortex might accurately encode differences in animacy (high variance), but only coarsely encode differences in color saturation (low variance).

For instance, if we were to show a human subject a set of object images that varied along the dimension of animacy (e.g., ranging from inanimate rocks to cats) we would expect them to clearly and accurately notice the differences between these images (see Figure B.1b). On the other hand, if we were to vary a more perceptually subtle property, such as color saturation, the differences might appear less pronounced and we could say that the visual system is less accurate along this dimension. At the extreme, we could generate images by sampling from a distribution of white noise, in which case differences would be almost imperceptible, despite varying significantly in the input space.

What these examples show is that the human brain does not accurately encode the vast space of all possible visual dimensions, but only a lower-dimensional subspace of these dimensions that are ecologically relevant for survival and behavior in the real world (i.e., dimensions along an *ecological subspace*). In the same way, any computational model of perception, such as a DNN, will preferentially encode different visual dimensions more or less accurately and define its own representational *model subspace*. We refer to dimensions along these representational subspaces as *latent*, and we refer to the dimensions of the larger visual space in which they are embedded as *ambient*.

With these concepts in mind, we can now begin to think about the conditions under which a model can achieve high encoding performance when predicting a given neural dataset. Intuitively, this can only happen when the stimuli span dimensions that are accurately encoded by *both* the model and the brain. In other words, the latent dimensions of the ecological subspace must overlap with latent dimensions of the model subspace. A key factor driving our simulated and empirical results is that the probability of these overlaps increases substantially as the latent dimensionality of the model subspace grows.

## Appendix C Implementation of simulations

Our process is summarized in Figure C.1, and consists of 3 steps. First, given our simulation parameters, we sampled subspace geometries within a high-dimensional ambient space. Next, we generated a set of experimental stimuli from the data subspace and projected them onto both the ecological and model subspaces, yielding a set of neural and model activations. Finally, we fit a linear encoding model to measure how well neural responses to the stimuli could be predicted from the corresponding model activations. By repeating this process across a range of simulation parameters (e.g., different model effective dimensionalities), we could better understand their influences on encoding performance. We describe this process in more detail below.

### Subspace geometry

In essence, our simulations consider four subspaces and relations between them: the natural image subspace, the data subspace from which experimental stimuli are sampled, the ecological subspace governing neural representations, and the model subspace. For simplicity, we parameterized all subspaces as multivariate Gaussian distributions embedded in a common ambient space of all possible visual dimensions. Using multivariate Gaussians also provided a simple method for modulating and measuring the subspace’s latent dimensionality, which we describe later in this section.

### Variance along latent dimensions

In our simulations, the amount of variance in a subspace along a given dimension has important significance. For the natural image and data subspaces, this variance corresponds to changes in a particular image feature (e.g., animacy). For the ecological and model representational subspaces, however, the variance along a dimension represents how accurately that dimension is encoded by the brain or a particular model. For example, consider Figure B.1b. Here, the human brain accurately and precisely represents differences in object animacy (high-variance dimension) but is relatively imprecise in how it represents fine changes in color saturation (low-variance dimension). One way to re-frame this idea is to think of the variance along a given dimension as representing its signal-to-noise-ratio (SNR).

### Sampling subspaces

Within an image space of ambient dimensionality *D_a_*, we wished to generate multivariate Gaussian subspaces with desired effective dimensionalities and mutual alignment pressures (Figure ^C^.^1^ step 1). First, we sampled a natural image subspace *_NI_* with effective dimensionality *ED_NI_*. The orthonormal eigenvectors for this subspace were sampled uniformly within the ambient dimensional space, whereas the eigenvalues were selected deterministically to achieve *ED_NI_*. Although there are many ways to design eigenspectra with a particular ED, we opted to parameterize the decay rate of the eigenvalues as a power law 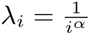 and solved for the *α* that yielded our desired ED.

We next sampled the ecological subspace 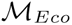, model subspace 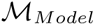, and data subspace 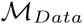. To select their eigenvalues, we followed the same power law parameterization as for the natural image subspace to achieve effective dimensionalities *ED_Eco_*, *ED_Model_*, and *ED_Data_*. Their eigenvectors, however, were all sampled in a way that depended on their respective alignment pressures to the natural image subspace 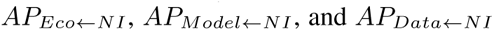. This aspect of the sampling procedure is described in detail below.

### Formulation of AP

In our simulations, AP is a scalar value that ranges between –1 and 1. An *AP* = 0 corresponds to no alignment pressure, in which case a basis of eigenvectors is sampled uniformly in the ambient space. When *AP >* 0, eigenvectors are sampled such that dimensions with larger eigenvalues capture more of the total variance in the natural image subspace (i.e., the high-variance dimensions of both subspaces are more likely to be aligned). On the other hand, when *AP <* 0, eigenvectors are sampled to preferentially align with low-variance dimensions of the natural image subspace.

Specifically, we needed to sample orthogonal vectors to form the eigenbasis of a new Gaussian subspace 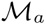, where those vectors preferentially spanned regions of high-variance in a reference Gaussian subspace 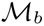, in a way that depended on *AP_a ← b_*. We achieved this by first defining a multivariate Gaussian distribution 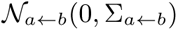. The eigenvectors of 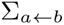 were equal to those of 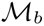, while the eigenvalues of 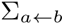 were generated as follows:

**Figure C.1:**
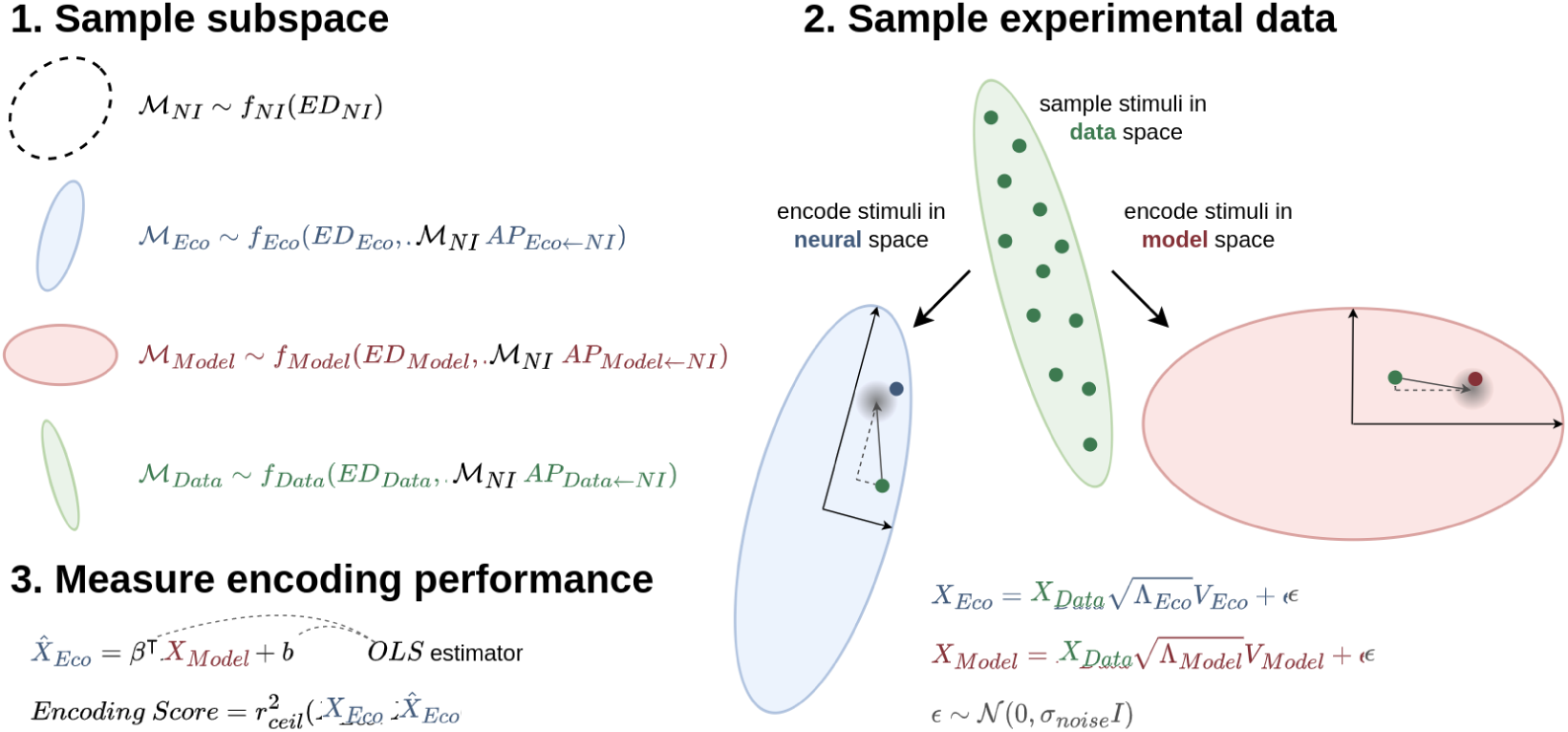
Simulating our theory of latent dimensionality and encoding performance. **1**. Our Gaussian subspaces were sampled to have a desired effective dimensionality. The ecological, model, and data subspaces were also sampled with a given alignment pressure to a shared space: the natural image subspace. **2.** Experimental data were sampled from the distribution specified by the experimental data subspace and then projected onto the ecological and model subspaces, which stretch or compress the data according to their variance along different dimensions. Isotropic noise was then added so that high-variance subspace dimensions had higher SNR and more accurately encoded their corresponding image features. **3.** A linear encoding model was trained using ordinary least squares (OLS) regression to predict neural responses from model activations. The reported encoding score is the percentage of explained variance normalized to the noise-ceiling of the neural data.

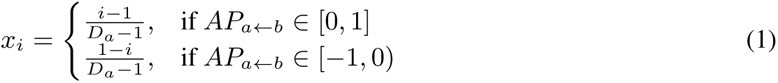

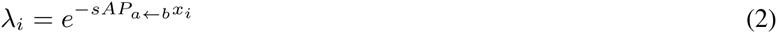

where *i* is the index of the eigenvalue starting from 1, *D_a_* is the ambient dimensionality of the subspaces, and *s* is a scaling factor which we set to 20. Essentially, this drew eigenvalues from *D_a_* equally spaced points on an exponential function in the domain of [0, 1] that is either decaying (in the case of positive AP) or growing (in the case of negative AP).

Next, we iteratively (1) sampled a vector 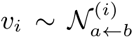, (2) normalized *v_i_* to unit length, and (3) projected 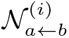 onto the subspace orthogonal to *v_i_*, giving 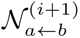. This process was repeated *D_a_* times. We then used the normalized *v_i_*’s as the eigenvectors of *M_a_*. First, note that the *v_i_*’s collectively define an orthonormal basis because each is sampled from a subspace orthogonal to 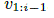. Second, note that for positive *AP_a ← b_* early *v_i_*’s are more likely to be oriented towards regions of high variance in *M_b_*, where 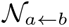 has most of its probability mass. For negative *AP_a ← b_*, though, the probability mass of 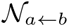 is concentrated along low-variance dimensions in *M_b_*, which results in early *v_i_*’s that tend to point in low-variance dimensions of *M_b_* as well. For an *AP_a ← b_* = 0, the covariance matrix of 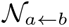 is identity, and the *v_i_*’s thus do not depend on the eigenvectors of *M_b_* in any way. Collectively, these properties satisfied our desiderata regarding the function of alignment pressure.

### Sampling experimental data

Having generated subspaces that specified the distribution of stimuli as well as model and ecological coding properties, our next step was to sample experimental data (Figure C.1 step 2). First, we sampled *N* points from the multivariate Gaussian distribution specified by the data subspace:

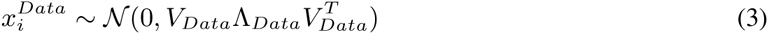

where *V_Data_* denotes the column matrix of data subspace eigenvectors and Λ*_Data_* is the diagonal eigenvalue matrix. These points can be thought of as experimental stimuli, which vary along different image dimensions.

Next, we projected the stimuli onto the eigenvectors of both the ecological and mode subspaces (*V_Eco_* and *V_Model_*) and scaled them by the standard deviation along each of those eigenvectors 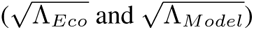. The net effect of this scaling was that the ecological/model subspaces amplified or attenuated different stimulus dimensions depending on whether or not they had significant variance along them. However, only applying this scaling would have had no effect on linear encoding performance, since regression weights could re-scale to compensate. Therefore, after performing this projection and scaling, we added ambient noise 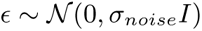 across all dimensions. The final result was a dataset of neural and model activations:

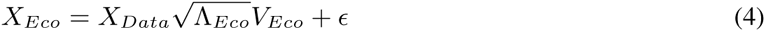

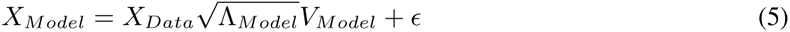

Since the magnitude of the noise was equal in all directions, its net effect was to modulate the SNR along the subspace dimensions. Essentially, ecological/model dimensions with high variance were relatively unaffected by the noise and accurately encoded stimulus features, whereas dimensions with low variance were dominated by the noise and only coarsely encoded stimulus features (e.g., the ordering of different stimuli along noise-dominated dimensions might not be preserved).

### Measuring encoding performance

After having stimulated neural and model activations, our last step was to measure the linear encoding performance of predicting *X_Eco_* from *X_Model_*. Specifically, we predicted neural activations 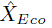 and then computed the percentage of explained variance normalized to the noise ceiling of *X_Eco_*:

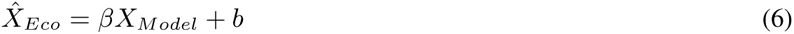

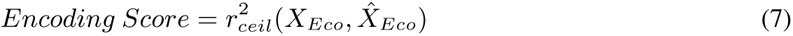

where the regression parameters *β* and *b* were estimated using ordinary least squares regression without any regularization and prediction accuracy was computed through cross-validation. The noise ceiling corresponded to the percentage of variance in *X_Eco_* that was explainable signal. In our simulations, we had direct access to this value because *E_Eco_* was generated according to:

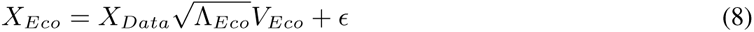

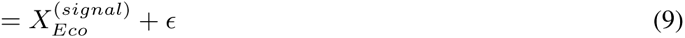

where 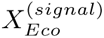 is the signal component of *X_Eco_*. Thus, we simply fit another linear regression model to predict *X_Eco_* using 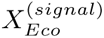 as regressors, in which case the resulting percentage of explained variance *r*^2^ corresponded to the noise ceiling.

When computing percentages of explained variance, we also needed to aggregate across all dimensions (i.e., neurons) of *X_Eco_* that were predicted. Typically, this is done by taking the mean 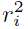 across all dimensions *i*, but this would violate an important principal of our theory wherein dimensions with larger variance contain more signal, and are therefore more important to predict. Instead, we computed a weighted average of all 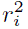, with weights equal to the variance in *X_Eco_* along dimension *i*.

### Simulation parameters

Unless otherwise stated, our simulation parameters were set as follows: *D_a_* = 100, *ED_NI_* = 20, *ED_Eco_* = 10, *ED_Data_* = 100, *AP_Eco←NI_* = 0.75, *AP_Model←NI_* = 0.75, *σ_noise_* = 0.1. *ED_Model_* ranged from 1 to *D_a_*, and 50 repeats of the simulation were performed for all values of *ED_Model_*, each with independently sampled subspaces/datasets.

## Appendix D Additional simulation results

In this section, we show how the relationship between *ED_Model_* and encoding performance can be modulated by different settings of *ED_Eco_*, *AP*, and *σ_noise_*. Figure D.1 demonstrates these effects, which we interpret below.

**Figure D.1:**
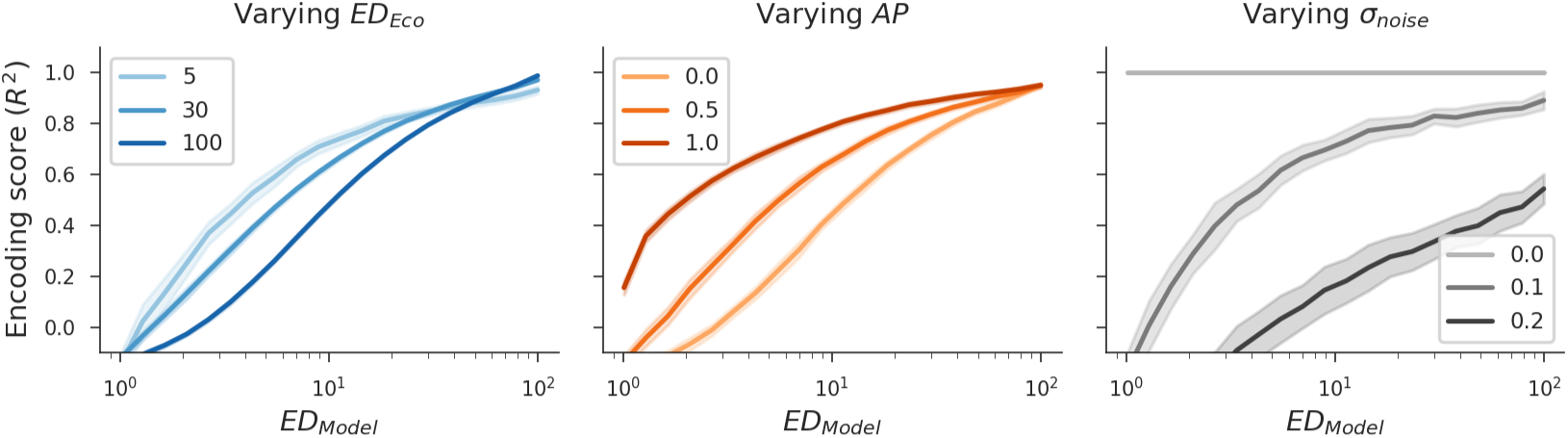
Modulating additional simulation parameters. Each simulation parameter modulated the relationship between *ED_Model_* and encoding performance. Within a plot, only the titled parameter was changed (shown in different line colors), while other parameters were held constant.

### Increasing

*ED_Eco_* As *ED_Eco_* increased, higher *ED_Model_* was needed to achieve the same level of encoding performance simply because there were more ecological dimensions to explain. In practice, this means that if the ecological subspace (i.e., representations in visual cortex) is high-dimensional, encoding performance will saturate later as a function of *ED_Model_*.

### Increasing

*AP* In regards to different *AP* between the model and ecological subspaces to the natural image subspace, we see that lower-dimensional models were able to achieve better encoding performance if they were preferentially aligned to ecological dimensions. Nevertheless, given any constant alignment pressure, there remains a positive correlation between *ED_Model_* and encoding performance, indicating independent contributions.

### Increasing

*σ_noise_* Varying *σ_noise_* is key to simulations of our theory. In the case of no noise (*σ_noise_* = 0), encoding performance was in fact independent of *ED_Model_*, and all models achieved perfect encoding performance. This is because our models had *some* non-zero variance along every ambient dimension, which, in the absence of noise, always led to an SNR of ∞. In essence, the variance along a dimension had no semantic meaning in this case because it could be scaled arbitrarily without any change in the representation. As *σ_noise_* increased, however, having high variance along a dimension became increasingly more important for accurately representing features with high SNR, and encoding performance therefore became more dependent on high *ED_Model_*.

### Limitations of our theory and simulations

While our simulations provide valuable intuitions regarding ED, AP, and encoding performance, they make several simplifying assumptions that are unlikely to hold in practice. First, they assume that all subspaces are multivariate Gaussians, in which case linear metrics such as ED are appropriate for estimating latent dimensionality. While the precise topologies of subspaces in biological and artificial neural representations are unknown, there is evidence that they are likely nonlinear [3]. Another simplification is that our simulations sample linear model and neural dimensions within the *same* ambient space, whereas in reality models and the brain both nonlinearly transform image dimensions. In other words, an image feature that is linearly encoded in the ecological subspace might be highly curved and warped in the model subspace. Future work could build on our simulation framework to explore these issues.

## Appendix E Details of DNN models

Table E.1 lists details for all DNNs used in our experiments. *PyTorch* models were obtained from the torchvision package and from PyTorch Hub [68], *VVS* models were obtained from Zhuang et al. [97], and *Taskonomy* models were obtained from Zamir et al. [94].

**Table E.1:** DNN models used in experiments.

| Training task | Learning setting | Training dataset | Architecture | Source |
| --- | --- | --- | --- | --- |
| Object classification | Supervised | ImageNet | AlexNet | PyTorch |
| Object classification | Supervised | ImageNet | VGG-16 | PyTorch |
| Object classification | Supervised | ImageNet | SqueezeNet | PyTorch |
| Object classification | Supervised | ImageNet | ResNet18 | PyTorch |
| Object classification | Supervised | ImageNet | ResNet50 | PyTorch |
| Barlow-Twins | Self-Supervised | ImageNet | ResNet50 | PyTorch |
| N/A | Untrained | N/A | AlexNet | N/A |
| N/A | Untrained | N/A | VGG-16 | N/A |
| N/A | Untrained | N/A | SqueezeNet | N/A |
| N/A | Untrained | N/A | ResNet18 | N/A |
| N/A | Untrained | N/A | ResNet50 | N/A |
| Object classification | Supervised | ImageNet | ResNet18 | VVS |
| Depth prediction | Supervised | ImageNet | ResNet18 | VVS |
| Auto-encoding | Self-supervised | ImageNet | ResNet18 | VVS |
| Colorization | Self-supervised | ImageNet | ResNet18 | VVS |
| Contrastive multiview coding | Self-supervised | ImageNet | ResNet18 | VVS |
| Contrastive predictive coding | Self-supervised | ImageNet | ResNet18 | VVS |
| Deep cluster | Self-supervised | ImageNet | ResNet18 | VVS |
| Instance recognition | Self-supervised | ImageNet | ResNet18 | VVS |
| Local aggregation | Self-supervised | ImageNet | ResNet18 | VVS |
| Relative position | Self-supervised | ImageNet | ResNet18 | VVS |
| SimCLR | Self-supervised | ImageNet | ResNet18 | VVS |
| Object classification | Supervised | Indoor buildings | ResNet50 | Taskonomy |
| Scene classification | Supervised | Indoor buildings | ResNet50 | Taskonomy |
| Semantic segmentation | Supervised | Indoor buildings | ResNet50 | Taskonomy |
| Curvature estimation | Supervised | Indoor buildings | ResNet50 | Taskonomy |
| Depth estimation | Supervised | Indoor buildings | ResNet50 | Taskonomy |
| Depth estimation (z-buffer) | Supervised | Indoor buildings | ResNet50 | Taskonomy |
| Edge detection (2D) | Supervised | Indoor buildings | ResNet50 | Taskonomy |
| Edge detection (3D) | Supervised | Indoor buildings | ResNet50 | Taskonomy |
| Egomotion | Supervised | Indoor buildings | ResNet50 | Taskonomy |
| Fixated pose estimation | Supervised | Indoor buildings | ResNet50 | Taskonomy |
| Non-fixated pose estimation | Supervised | Indoor buildings | ResNet50 | Taskonomy |
| Keypoint detection (2D) | Supervised | Indoor buildings | ResNet50 | Taskonomy |
| Keypoint detection (3D) | Supervised | Indoor buildings | ResNet50 | Taskonomy |
| Point matching | Supervised | Indoor buildings | ResNet50 | Taskonomy |
| Reshading | Supervised | Indoor buildings | ResNet50 | Taskonomy |
| Room layout estimation | Supervised | Indoor buildings | ResNet50 | Taskonomy |
| Surface normal estimation | Supervised | Indoor buildings | ResNet50 | Taskonomy |
| Vanishing point estimation | Supervised | Indoor buildings | ResNet50 | Taskonomy |
| Auto-encoding | Self-supervised | Indoor buildings | ResNet50 | Taskonomy |
| Denoising | Self-supervised | Indoor buildings | ResNet50 | Taskonomy |
| Inpainting | Self-supervised | Indoor buildings | ResNet50 | Taskonomy |
| Jigsaw | Self-supervised | Indoor buildings | ResNet50 | Taskonomy |
| Unsupervised segmentation (2D) | Self-supervised | Indoor buildings | ResNet50 | Taskonomy |
| Unsupervised segmentation (2.5D) | Self-supervised | Indoor buildings | ResNet50 | Taskonomy |

## Appendix F Additional analyses of ED and encoding performance

Here, we replicate our main results in more settings. Figure F.1 shows the relationship between effective dimensionality and encoding performance without applying an average-pooling operation to the DNN feature maps. Figure F.2 colors models according to what dataset they were pre-trained on (ImageNet or Taskonomy). Figures F.3 and F.4 show encoding performance across multiple species, recording modalities, and brain regions. Figure F.5 fits encoding models using OLS instead of partial-least-squares regression. Figure F.6 shows results for only the layer of each model that achieved the highest encoding performance, rather than all layers.

Our general results hold across all of these settings, with the exception of V1 in the monkey electrophysiology data. We speculate that this is due to the far lower complexity of representations in V1, which serve primarily as simple edge detectors for later processing in higher-level regions.

**Figure F.1:**
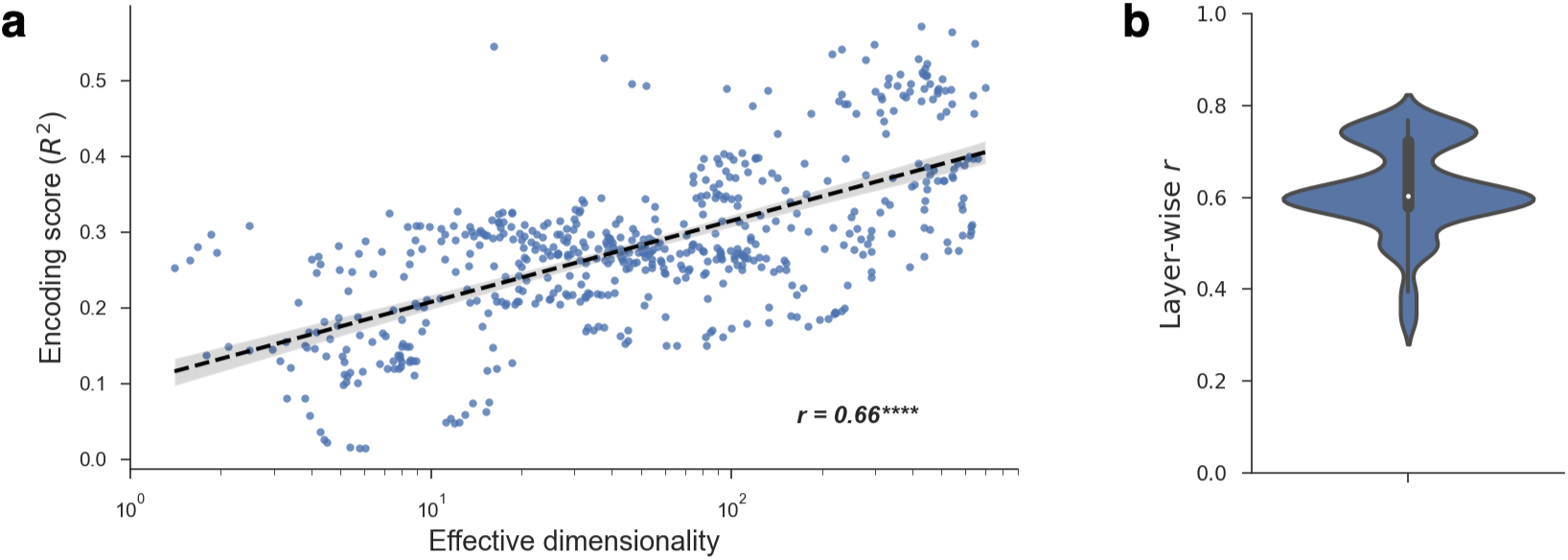
Effective dimensionality and encoding performance without average-pooling. **a**. The encoding performance achieved by a model scaled with the effective dimensionality of its entire feature map (without average-pooling applied). Each point in the plot was obtained from one layer from one DNN, resulting in a total of 568 models (see main text for further details). **b.** Even when conditioning on a particular DNN layer, controlling for both depth and ambient dimensionality, effective dimensionality and encoding performance continued to strongly correlate. The plot shows the distribution of these correlations (Pearson *r*) across all unique layers in our analyses.

**Figure F.2:**
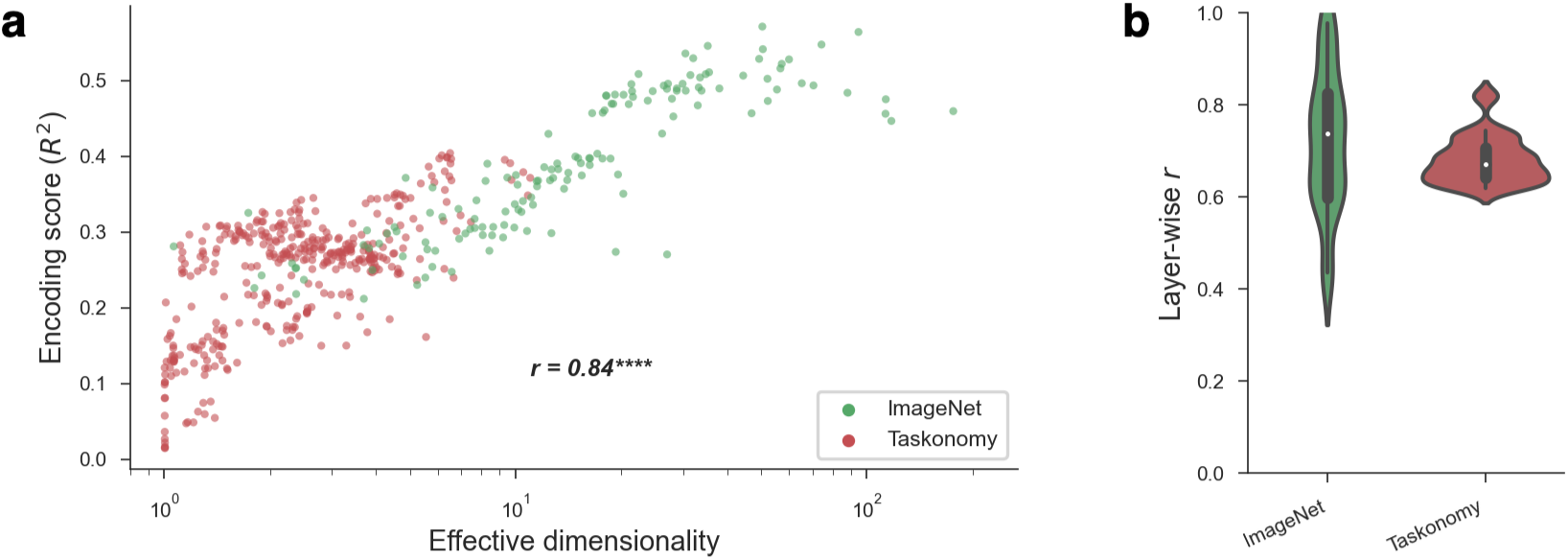
Effective dimensionality and encoding performance by training set. **a**. The encoding performance achieved by a model scaled with the effective dimensionality of its features, within both ImageNet- and Taskonomy-trained models. **b.** Even when conditioning on a particular DNN layer, controlling for both depth and ambient dimensionality, effective dimensionality and encoding performance continued to strongly correlate for both training sets. The plot shows the distribution of these correlations (Pearson *r*) across all unique layers in our analyses.

## Appendix G ED and Representational Similarity Analysis

To provide additional evidence that our central results are not due to trivial statistical effects wherein models with higher latent dimensionality have more degrees of freedom to predict neural data, we replicated our results using representational similarity analysis (RSA) [53]. In RSA, a dissimilarity matrix is constructed for both the model and the brain data by computing a distance between the representations for each pair of stimuli. These matrices are then correlated to evaluate their similarity. Importantly, unlike when fitting encoding models, this method for measuring the similarity between a model and the brain is entirely non-parametric and thus cannot be biased to favour models with high latent dimensionality.

**Figure F.3:**
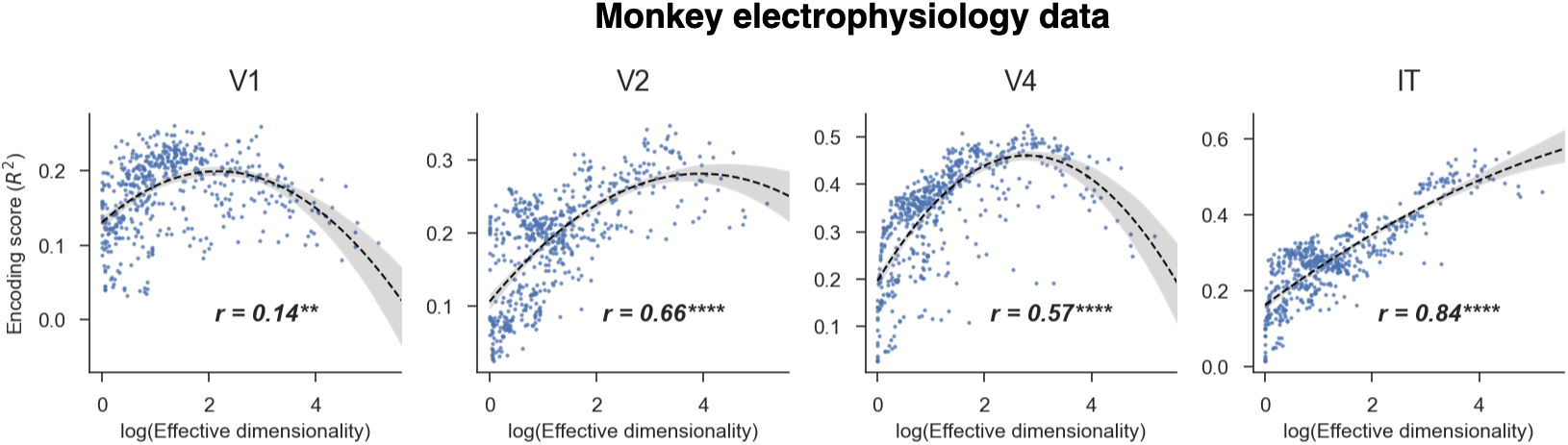
Latent dimensionality and encoding performance on monkey electrophysiology data. The encoding performance for all of our models across multiple brain regions in monkey electrophysiology datasets collected by Majaj et al. [62] (IT and V4) and Freeman et al. [28] (V1 and V2), plotted against the models’ ED. Our results hold across all brain regions except for V1; encoding performance increases with latent dimensionality. Another possible interpretation of these results, which becomes apparent when fitting inverted-U curves to the data (2nd-order polynomial regression), is that each region (including V1) has an “optimal” model ED reminiscent of the “Joint regime” illustrated in Figure 1c, which tends to increase along the cortical hierarchy. Note that optimal model ED values can only be compared within (V1, V2) and (V4, IT), since recordings for each set of regions were obtained from different stimulus sets.

**Figure F.4:**
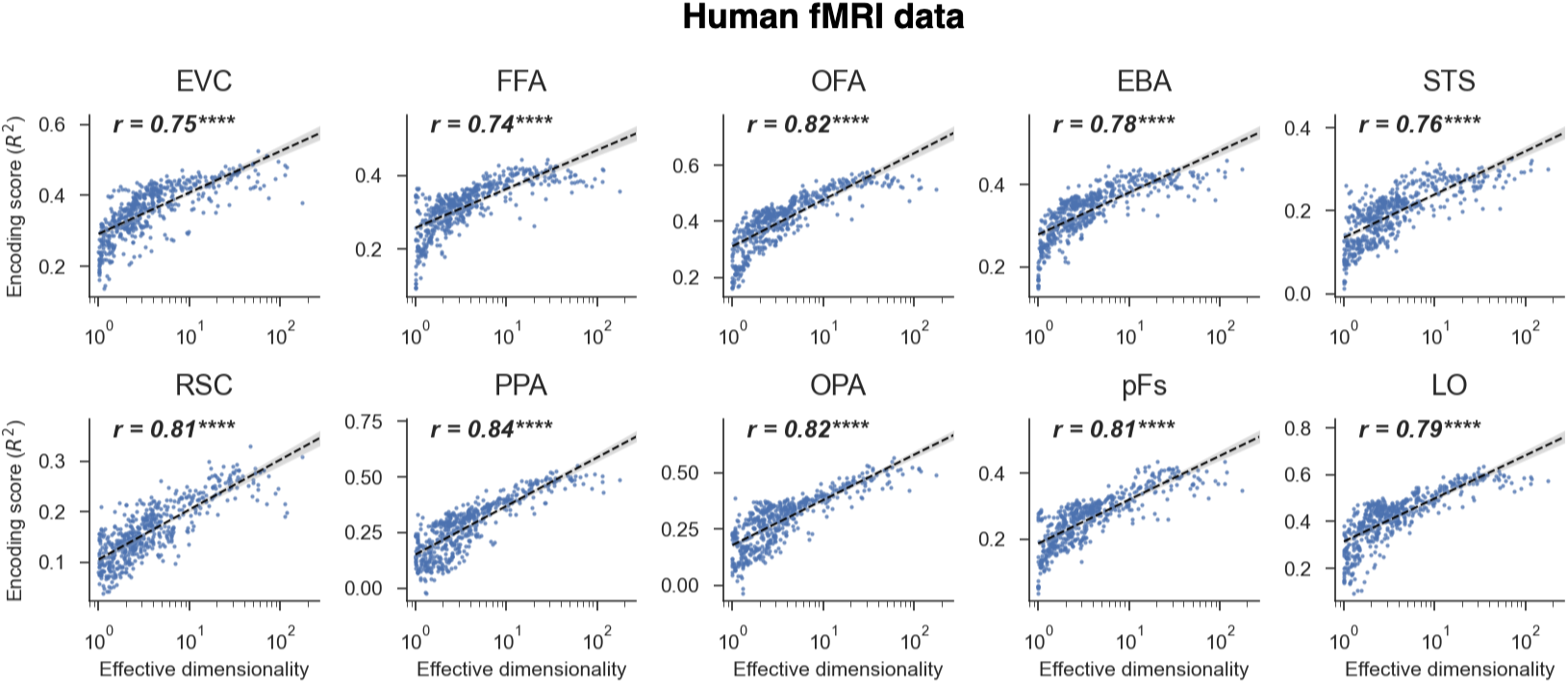
Latent dimensionality and encoding performance on human fMRI data. The encoding performance for all of our models across multiple brain regions in a human fMRI dataset collected by Bonner and Epstein [9], plotted against the models’ ED. Our results hold across all brain regions; encoding performance increases with latent dimensionality. EVC=early visual cotex, FFA=fusiform face area, OFA=occipital face area, EBA=extrastriate body area, RSC=retrosplenial complex, PPA=parahippocampal place area, OPA=occipital place area, STS-superior temporal sulcus, LO=lateral occipital region, pFs=posterior fusiform region.

While not as strong, there is nevertheless a clear trend in which models with higher RSA scores tend to have higher ED. Thus, our core results replicate using RSA.

## Appendix H ED varies with model and training parameters

To better understand how and why latent dimensionality varies in DNNs (and perhaps manipulate it for a desired effect), we can start by observing its empirical relationship to parameters of the training procedure, dataset, and architecture. Figure H.1 illustrates several of these relationships, which we were able to quantify thanks to the use of our large bank of models. We summarize our most important conclusions from these analyses below.

**Figure F.5:**
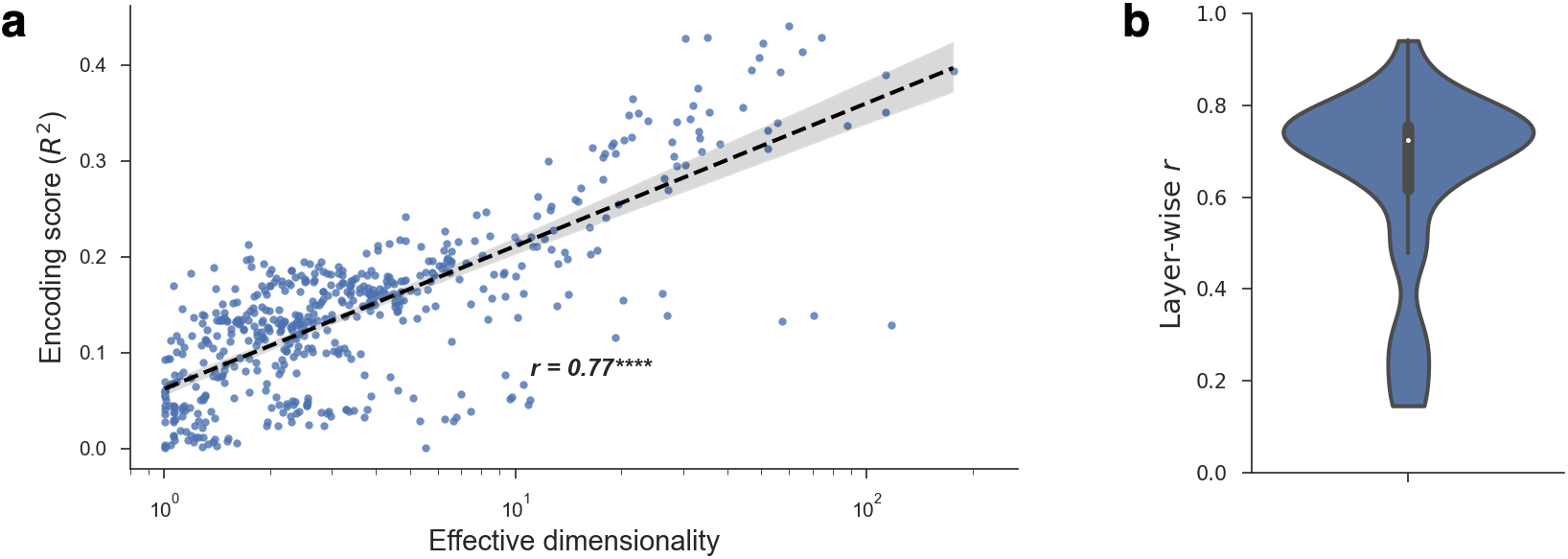
Effective dimensionality and encoding performance using OLS regression. **a**. The encoding performance achieved by a model fit using OLS regression scaled with its effective dimensionality. Each point in the plot was obtained from one layer from one DNN, resulting in a total of 568 models (see main text for further details). **b.** Even when conditioning on a particular DNN layer, controlling for both depth and ambient dimensionality, effective dimensionality and encoding performance continued to strongly correlate. The plot shows the distribution of these correlations (Pearson *r*) across all unique layers in our analyses.

**Figure F.6:**
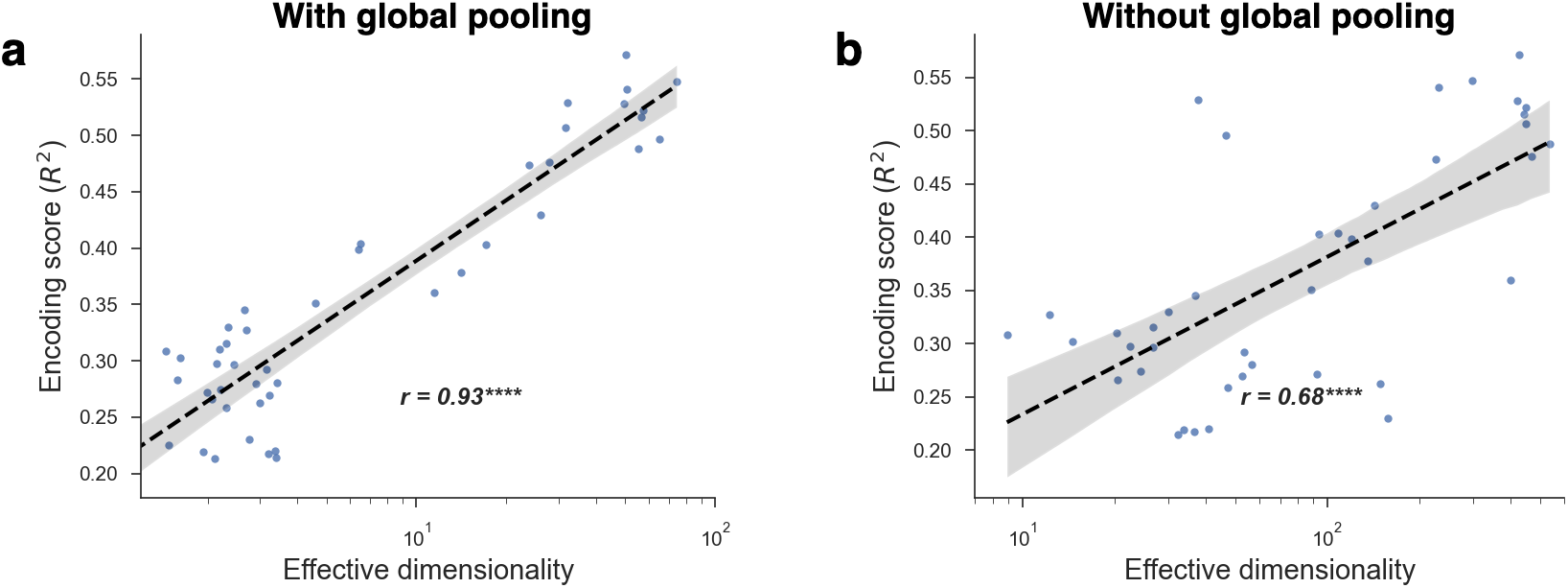
Effective dimensionality and encoding performance using the best layer from each model. Instead of showing results for all layers of each DNN, an alternative method is to consider only the layer that achieves the best encoding performance as the “model”. Even using this approach, there is still a clear trend between model ED and encoding performance. **a.** With global pooling applied to the model features prior to computing ED. **b.** Without global pooling applied.

### Training increases effective dimensionality

Figure H.1a shows how effective dimensionality varied across the layer hierarchy for multiple CNN architectures when they were trained on ImageNet object classification compared to when they were untrained and had randomly initialized weights. We can see that training resulted in substantial increases in effective dimensionality for both architectures across all layers. To solve complex tasks such as object classification, then, it appears that models must learn to extract a large number of orthogonal image features. This finding contradicts a commonly held belief that DNNs trained on visual tasks compress high-dimensional inputs to a small number of latent dimensions [72, 17, 49, 3, 23].

### Effective dimensionality increases with layer depth

Another notable trend in Figure H.1a is that effective dimensionality increased as a function of layer depth within the two supervised classification models we considered. Importantly, this cannot be explained simply as a result of an increasing number of channels along the layer hierarchy, as effective dimensionality remained more or less constant within the untrained models. This gradual increase in effective dimensionality appears to contradict other findings from [3, 19, 17] in which latent dimensionality generally decreases as a function of layer depth, but there are important methodological differences to note. First and foremost, we computed effective dimensionality only along the channel dimension of our feature maps after applying a average-pooling operation across the spatial dimensions, which allowed us to focus on the diversity of image features. Given that spatial resolution decreases as a function of layer depth in our architectures (and most convolutional DNNs in general), the effective dimensionality of earlier layers will be higher simply due of the larger number of spatial dimensions that they contain. Another important difference in our work is that we only considered convolutional layers and performed no analyses on fully-connected layers. Indeed, much of the drop in latent dimensionality reported in other work occurs within these fully-connected layers.

**Figure G.1:**
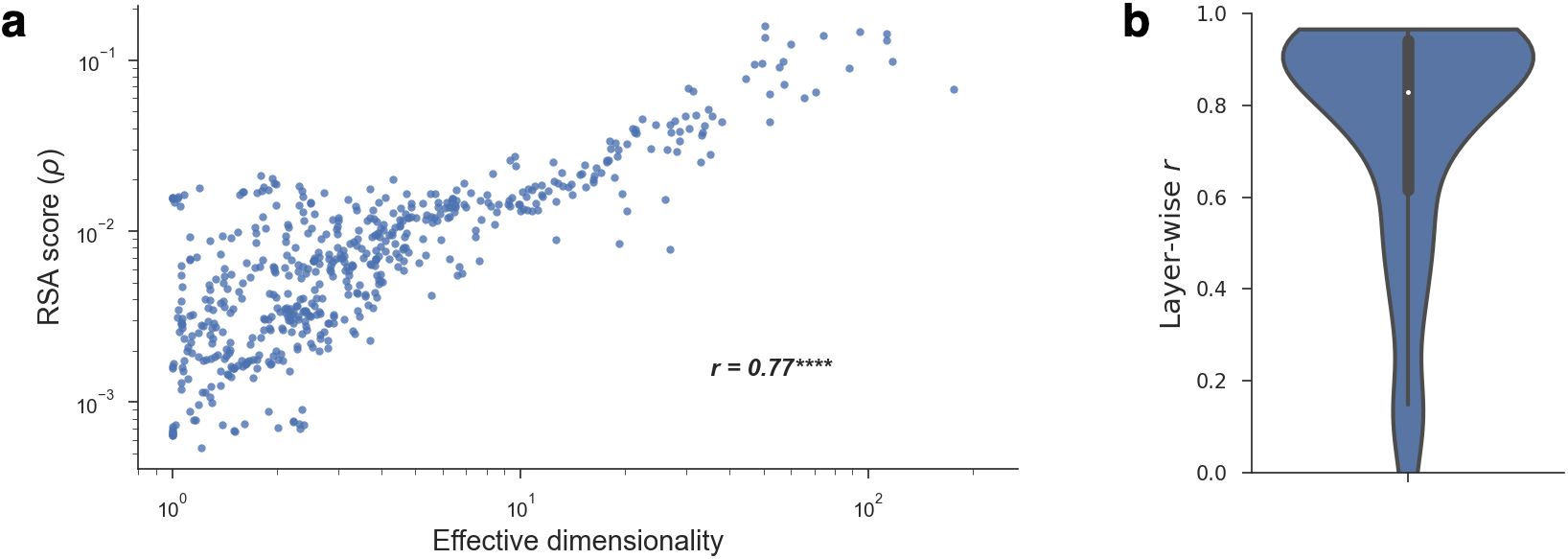
Effective dimensionality and Representational Similarity Analysis (RSA). We compared the representations in our models to those in the monkey IT electrophysiology data using RSA. Note that the y-axis is on a log-scale in order to provide better resolution in face of the high variation in RSA scores. Our results hold across this different similarity metric; the similarity between model and brain representational dissimilarity matrices increases with latent dimensionality.

**Figure H.1:**
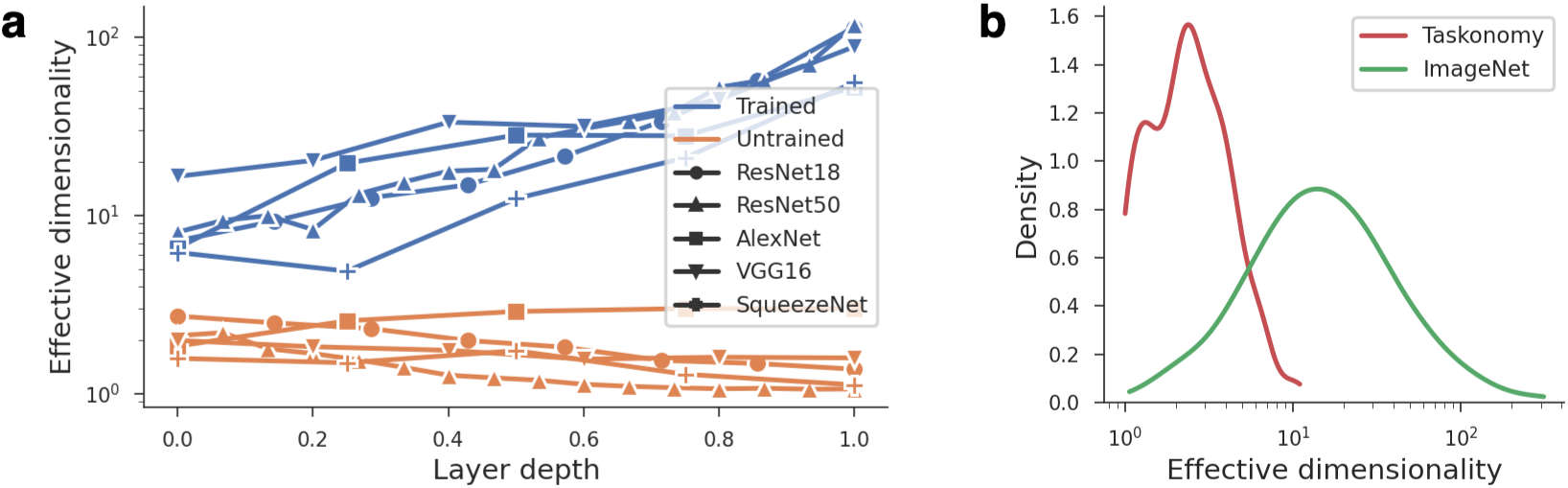
Effective dimensionality varies with model and training parameters. **a**. Models trained on object classification (blue) had larger effective dimensionality than untrained models (orange) across layers in multiple architectures. After training, effective dimensionality also gradually increased as a function of layer depth (only convolutional layers are shown). **b.** Plots indicate the distribution of effective dimensionality across models that were trained on Taskonomy (red) and ImageNet (green). These distributions differed significantly, despite the models in both groups largely sharing similar architectures and training tasks.

### Training data has a large impact on effective dimensionality

Another important factor that has a significant impact on a model’s learned representations is the training dataset. Our bank of models includes DNNs trained on ImageNet and Taskonomy. Figure H.1b shows the distribution of effective dimensionality for all models trained on each of these datasets. Despite similar architectures and training tasks used on both datasets, the ImageNet-trained models tended to have significantly larger effective dimensionality. Although we did not perform further analyses to determine which dataset differences explain this result, we speculate that it is due to the much greater diversity of image statistics within ImageNet. Whereas ImageNet contains images spanning many object categories appearing in diverse environments, Taskonomy consists solely of man-made indoor scenes across 600 buildings. Effective dimensionality, therefore, might scale in proportion to the complexity and variation of image features in the training data.

## Appendix I High ED alone is not sufficient to yield strong performance

Our findings show that ED is positively correlated with encoding performance when examining standard DNNs used in computational neuroscience. However, it is important to emphasize that high ED alone is not sufficient to yield strong encoding performance. Indeed, it is not difficult to imagine contrived models with extremely high dimensionality but no predictive power. As a simple example, imagine a maximally sparse representation in which each stimulus elicits a response along a single, unique dimension (akin to “grandmother cells” [5]). In this case, because every stimulus is represented along a unique dimension, an encoding model fit to a training set would have no ability to generalize to unseen stimuli.

In this section, we discuss some of the necessary conditions for observing a strong and positive correlation between effective dimensionality and encoding performance. In addition, we provide some empirical experiments to support our arguments.

### Alignment pressure plays a significant independent role

In the vocabulary of our theory laid out in Section 2.1, encoding performance depends not only on the latent dimensionality of a model, but also on its alignment pressure. And, because latent dimensionality can vary independently from alignment pressure (in the sense of independent causal interventions), it is not causally sufficient for achieving good encoding performance. Furthermore, in the infinite-dimensional space of possible visual features where model dimensions are unlikely to overlap with ecologically-relevant ones by chance, alignment pressure is essential.

In empirical models of visual cortex, the statistical relationship between alignment pressure and latent dimensionality has not been investigated. If models achieving high alignment pressure to biological representations tend to systematically have lower latent dimensionality, we might observe a net negative correlation between latent dimensionality and encoding performance. That this is not the case in our empirical results suggests that alignment pressure is probably uncorrelated (or perhaps positively) correlated with latent dimensionality.

### Latent dimensionality must reflect the number of accurately-encoded features

If high latent dimensionality improves encoding performance in all circumstances, we should be able to trivially obtain excellent encoding performance with any model by applying simple feature transformations. ZCA whitening, for instance, applies a linear transformation that results in a new set of features with covariance matrix close to identity (i.e., a flat eigenspectrum with maximum effective dimensionality). After applying ZCA to all models, however, we saw no improvement in encoding performance, despite significant increases in effective dimensionality, as shown in Figure I.1. Again, this shows that effective dimensionality alone is not sufficient to improve encoding performance. In this case, the reason is that ZCA whitening does not augment the model with additional information about the stimulus. Our theory in Section 2.1 states that the relationship between latent dimensionality and encoding performance depends on an assumption that higher-variance dimensions more accurately encode stimulus features. ZCA whitening, however, violates this assumption since it increases latent dimensionality by numerically scaling existing model dimensions without changing their semantics. No new dimensions are added and no existing dimensions are encoded more accurately, so encoding performance remains unchanged.

**Figure I.1:**
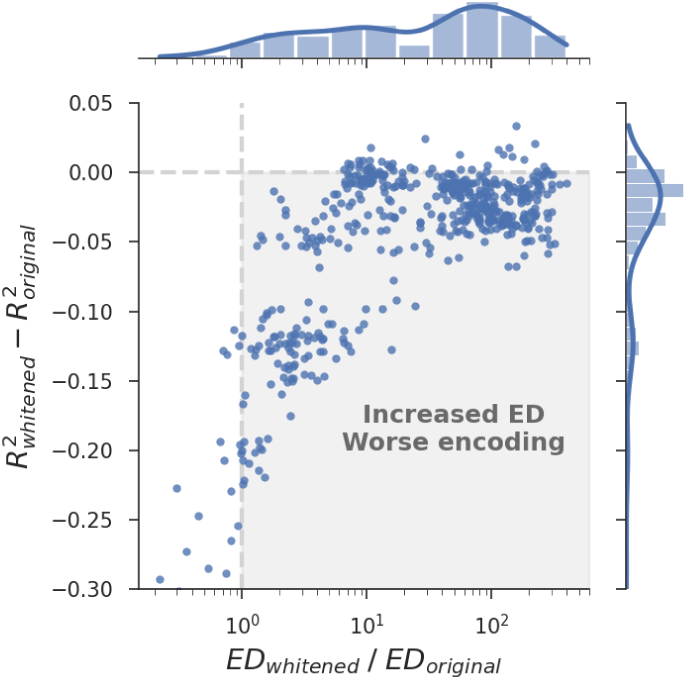
Increasing latent dimensionality with ZCA whitening does not enhance encoding performance. The y-axis shows the difference in encoding performance after whitening model features, while the x-axis shows the ratio of increase in effective dimensionality. Most whitened models saw a substantial increase in effective dimensionality, but showed either no change in encoding performance or a decrease (highlighted gray region).

### The empirical relationship between dimensionality and encoding performance is robust

Despite the above caveats, we note that it is difficult to construct poor-performing high-dimensional models in practice, without having to resort to trivial feature transformations such as whitening. We attempted to do so by training a DNN on a version of ImageNet where the labels in the training set were randomly scrambled. Due to their large capacity, it is well known that DNNs are able to achieve low training error on this task by finding an arbitrary mapping between each input and its label, essentially memorizing the dataset [95]. Our rationale for choosing this task was that it is unlikely to produce ecologically-relevant dimensions, but stands a good chance of learning a high-dimensional latent space in which it is easier to linearly separate arbitrarily labeled data [40]. However, this turned out *not* to be the case. In Figure I.2, we show the effective dimensionality and encoding performance of a DNN fit to scrambled labels and compare it to an identical architecture fit with the correct labels. As expected, the DNN with scrambled labels achieved much lower encoding performance. Surprisingly, however, this model also had much lower effective dimensionality than its correctly trained counterpart. We speculate that ecologically-relevant visual tasks in which humans excel (and most DNNs are trained on) require high latent dimensionality as a result of their inherent complexity, producing a positive correlation between latent dimensionality and alignment pressure. We explore this possibility in Section 2.4.

**Figure I.2:**
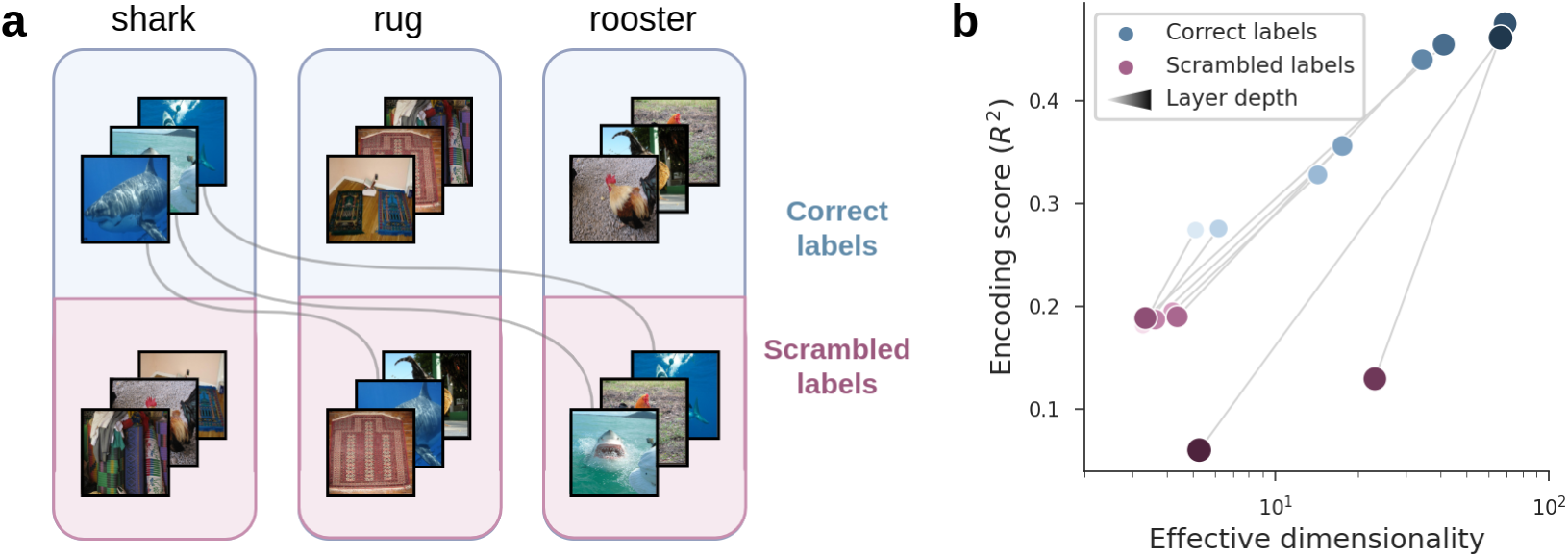
Training a model to overfit scrambled labels does not increase latent dimensionality. **a**. We trained the same ResNet18 DNN architecture on ImageNet classification in two different settings: once with correctly labeled images (blue) and the other time with scrambled labels, such that each image was assigned to a random class. Despite the arbitrary nature of the second task, the model was able to achieve good performance on the training set (47% accuracy over 1000 classes) by memorizing an idiosyncratic mapping from each input to its label. **b.** Our initial hypothesis was that the model trained with scrambled labels would have higher effective dimensionality and lower encoding performance than the model trained with correct labels, but our results surprisingly run counter to this intuition: the model trained with scrambled labels had lower encoding performance and *lower* effective dimensionality. Blue points denote layers from the model trained with correct labels, and purple points denote layers from the model with scrambled labels. Size and brightness denote increasing layer depth, and lines indicate matching layers.

## Appendix J Other correlates of DNN encoding model performance

In this section, we directly address alternate hypotheses proposed in the literature to explain the performance of DNN models of visual cortex. While these can be cast as competing hypotheses to our proposed theory based on ED, we will argue that task-performance measures and geometric properties should be instead viewed as different levels of explanation that provide complementary views of the function and underlying structure of neural representations.

### Object classification performance

A commonly held view is that DNNs explain neural data because their representations are optimized to perform well on ethologically-relevant tasks, such as object classification [93, 92, 48]. Indeed, in our models, we found that few-shot classification performance on object categories from ImageNet-21k predicted encoding performance very well (Figure J.1a). However, it is important to note that task performance and ED are not competing theories but, rather, different levels of explanation that provide complementary insights into the nature of neural network representations. Models that perform ethologically-relevant visual tasks are likely to explain neural responses, because the brain is optimized for such tasks as well and the space of good solutions is likely limited. However, this does not tell us *how* the models solve these tasks, for which we must look at the underlying representational structure. In the language of Marr [63], theories based on classification performance function at the abstract computational level, but there must also be explanations for DNN encoding performance that function at the representational level. ED is one such representation-based explanation. In Sections 2.4 and 2.5, we showed how ED empirically predicts classification performance on novel categories and provided one theoretical explanation for why this happens, which was proposed by Sorscher et al. [85]. Furthermore, when we conducted a variance partitioning analysis [65, 8] of encoding performance as a function of both ED and classification performance (Figure J.1b), we found that nearly all of the variance was shared. In sum, there is good reason to believe that ED and classification performance (or visual task performance more generally) are causally intertwined, and that they jointly provide a richer explanation of DNN models of visual cortex.

### Other geometric properties of the representations

To compare our theory to other representation-based hypotheses, we considered several geometric properties of representations that were described in Sorscher et al. [85] to predict the few-shot classification performance of pre-trained DNNs. For instance, one such geometric property is *signal*, which quantifies the average distance between pairs of class centroids in the representation. Some of these properties also relate to ED, except that they are computed either per object category and then averaged (“within-concept ED”), or they are computed using object category centroids (“between-concept ED”). Finally, the signal-to-noise ratio property (SNR) is an aggregate metric that combines all of the others to predict few-shot classification performance on novel categories using a prototype learning rule. See Sorscher et al. [85] and their related code (or ours) for a more complete description of these geometric properties and how they were computed. We used the same setup, with the same ImageNet-21k categories as Sorscher et al. The one exception is “between-concept ED”, which was not computed in Sorscher et al. [85], and simply consists of computing ED using object category centroids as the samples. The relationships between all of these geometric properties and encoding performance are shown in Figure J.1c.

We found that ED-related properties of the representation (i.e., global, within-concept, and between-concept ED) have the strongest relationship to encoding performance, with the exception of the aggregate SNR metric. This shows that among several geometric statistics of a representation, a simple measure of its latent dimensionality can most accurately predict its performance as a model of higher visual cortex. We wondered whether even the performance of SNR—a more complex aggregate metric that incorporates ED—could largely be accounted for by ED as well. We therefore conducted variance partitioning analyses of both encoding performance and classification performance on ImageNet-21k using ED and SNR as the predictors (Figure J.1d). Indeed, we found that almost all of the variance is shared, which suggests that ED is the primary driver of high SNR. We then conducted similar variance partitioning analyses using ED and “signal” as the predictors, since “signal” was the metric that best predicted encoding performance among non-ED metrics. Surprisingly, we found that ED explained significant unique variance, while “signal” did not. It therefore appears that “signal” only explains encoding and classification performance insofar as it correlates with ED, whereas ED explains unique variance in model performance.

**Figure J.1:**
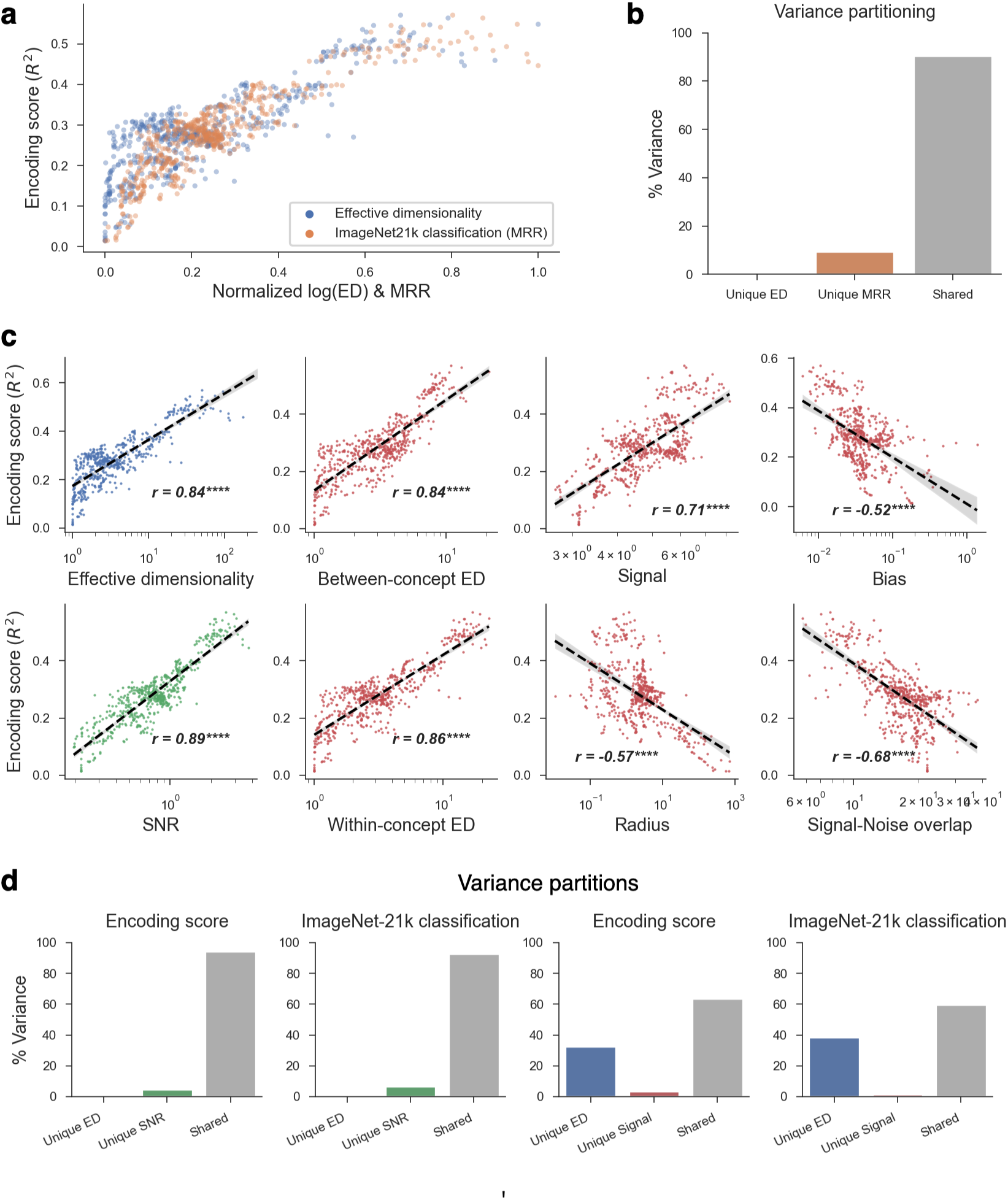
A theory of encoding performance based on ED is compatible with one based on classification performance, and superior to one based on other geometric properties of a representation. **a**. Encoding performance vs. ED (blue) and ImageNet-21k classification performance quantified using the mean reciprocal rank (MRR, orange). Both metrics were highly predictive of encoding performance. Note that the x-axis shows both metrics normalized to be between (0-1) so that they can easily be compared in the same plot. **b.** A variance partitioning analysis showed that ED and classification performance explain almost no unique variance in encoding performance, with about 90% of it being shared. This suggests that ED and classification performance should not be considered as independent and competing hypotheses, but are rather different levels of explanation that provide a complementary perspective. **c.** Encoding performance vs. ED (blue) and other geometric properties of a representation introduced in Sorscher et al. [85] (red for most properties, green for SNR—an aggregate property computed from the others). Aside from SNR, ED-related metrics had the strongest relationship to encoding performance. **d.** We partitioned the variance explained in both encoding performance and classification performance by ED, SNR, and “signal”, which is another key geometric property. SNR shares almost all of its variance with ED, while ED explains unique variance. All encoding performance scores are for the monkey IT electrophysiology data.

